# Genome-Wide Transcriptomics Uncovers a Lung-to-Aorta Inflammatory Program Driving Influenza-Induced Atherosclerosis Exacerbation

**DOI:** 10.64898/2026.10.22.761022

**Authors:** Louis DeVito, John C. Notaro, Nicholas J. Constantinesco, Dhivyaa Rajasundaram, John F. Alcorn, Radha Gopal

**Affiliations:** Department of Pediatrics, University of Pittsburgh, PA, United States

**Author notes:** Correspondence: Radha Gopal One Children’s Hospital Dr. 9129 Rango’s Research Center, 4401 Penn Ave. Pittsburgh, PA 15224.

## Abstract

Influenza infection increases cardiovascular risk in individuals with pre-existing atherosclerosis, yet the genome-wide transcriptional basis of this association remains undefined. We performed bulk RNA sequencing of aorta and lung tissues from high-fat diet-fed *Apoe*−/− mice infected with influenza A/PR/8/34 versus PBS-treated controls at day 15 post-infection, identifying 6,828 differentially expressed genes across both tissues (5,694 in the lung and 1,134 in the aorta; adjusted p < 0.05). In the lung, influenza sustained a CXCR3-driven chemokine storm, comprehensive Th1, Th17, and exhaustion T-cell activation programs, coordinated monocyte recruitment through *Ccl2*, *Ccl7*, *Ccl8*, and *Ccl12*, a broad acute phase response including *Saa3* and *A2m*, alveolar identity collapse through loss of *Nkx2-1* and surfactant proteins, and broad metabolic dysregulation involving suppression of fatty acid oxidation, mitochondrial function, and circadian metabolic regulators. In the aorta, influenza drove macrophage and DAMP activation through *Trem1*, *Fpr2*, and *S100a8*, cytotoxic T-cell engagement, smooth muscle cell dedifferentiation through *Myocd* loss, convergent bioenergetic failure, and an ectopic lung-identity gene signature representing a previously undescribed cross-organ communication mechanism. Circadian clock and Hippo/YAP pathway suppression were shared across both tissues, identifying systemic vulnerabilities connecting pulmonary infection to vascular structural compromise. Consistent with the in vivo findings, influenza-infected *Apoe*−/− bone marrow-derived macrophages exhibited inflammasome activation, DAMP amplification, M1 polarization, and suppression of prostaglandin-mediated inflammatory resolution. Together these findings define a lung-to-aorta inflammatory and metabolic axis driving atherosclerotic plaque destabilization during the post-infection recovery period and identify CCR2/CXCR3 antagonism, NLRP3 inhibition, and YAP pathway activation as candidate therapeutic interventions.

**Graphical abstract:** Influenza infection drives atherosclerotic plaque destabilization

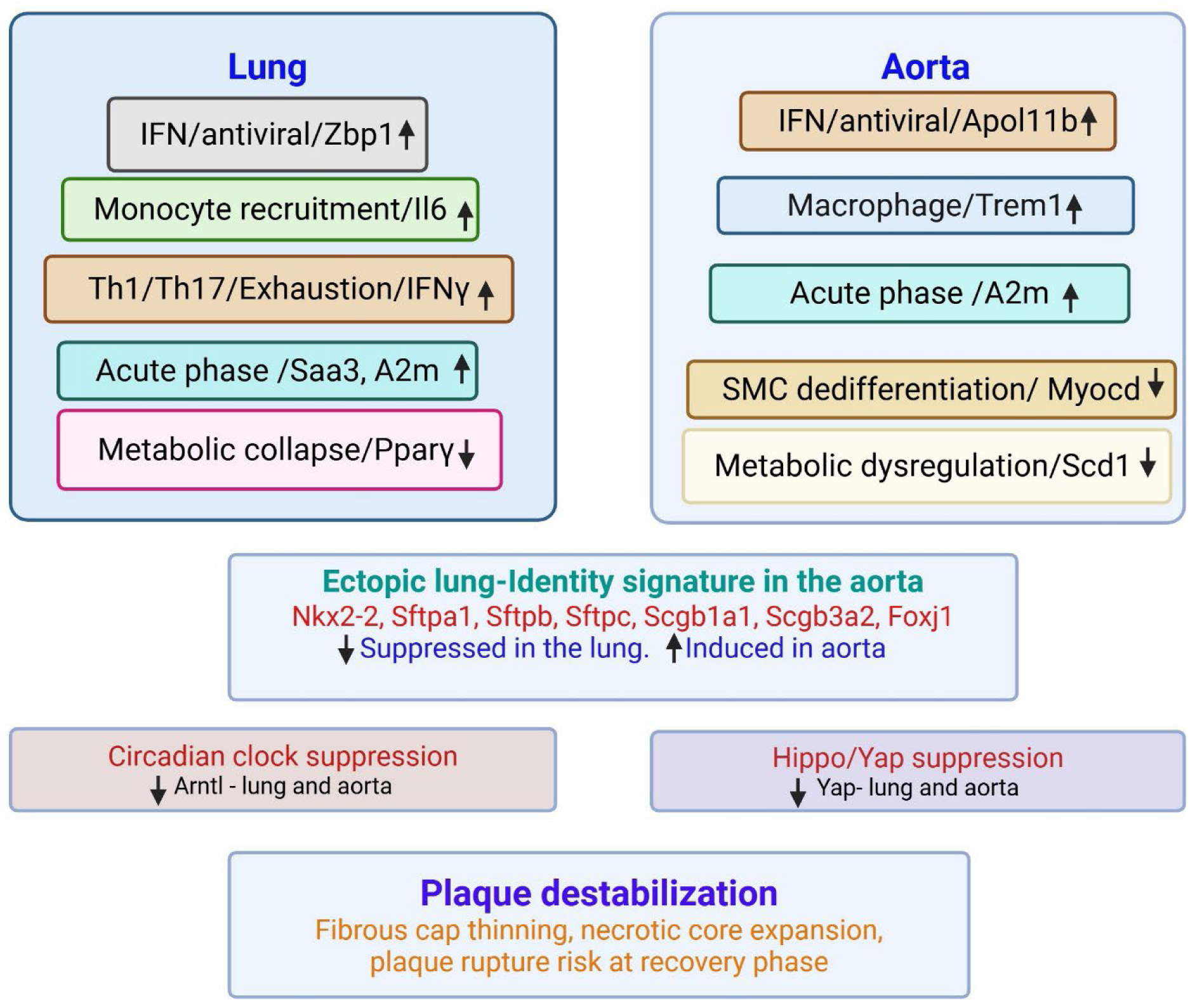

## 1. Introduction

Atherosclerosis is a chronic inflammatory disease of the arterial wall characterized by the accumulation of lipid-laden plaques, driven by dysregulated lipid metabolism, immune activation, and vascular endothelial dysfunction (1, 2). Plaque rupture, the precipitating event in most myocardial infarctions and strokes, results from a shift in the plaque microenvironment from a stable, smooth muscle-rich state to an unstable, inflammation-dominated state characterized by macrophage infiltration, matrix metalloproteinase-mediated fibrous cap thinning, and smooth muscle cell phenotypic switching (1–5). While the pathogenesis of chronic atherosclerosis is well defined, the molecular mechanisms by which acute infectious triggers accelerate this transition remain incompletely understood.

Epidemiological evidence strongly implicates influenza infection as a cardiovascular risk amplifier. Influenza seasons correlate consistently with increased hospitalizations and deaths from cardiovascular disease (6–8). Laboratory-confirmed influenza infection increases the risk of myocardial infarction six-fold during the acute phase, and the 2009 H1N1 pandemic alone was estimated to cause 83,300 cardiovascular deaths globally (9, 10). Influenza drives cardiovascular pathology through multiple mechanisms. Direct viral infection of vascular endothelial cells induces pro-inflammatory chemokines, adhesion molecules, and matrix metalloproteinases. Systemic IFN-γ production promotes macrophage foam cell formation and vascular smooth muscle cell synthetic phenotype switching. Innate immune activation further amplifies plaque inflammation (3, 11, 12). Critically, cardiovascular risk remains elevated for weeks after clinical recovery, suggesting that the transcriptional consequences of infection persist well beyond the acute phase (6–8). Despite this cellular and protein-level understanding, the genome-wide transcriptional basis of this persistent cardiovascular risk has not been characterized.

The *Apoe*−/− mouse model fed a high-fat diet is the most widely used system for studying accelerated atherosclerosis (13). Deficiency of apolipoprotein E (Apoe) leads to impaired lipoprotein clearance and rapid plaque formation, and *Apoe*−/− cells show enhanced influenza virus binding, making this model particularly relevant to the influenza-atherosclerosis interaction (13). Previous studies in this model have established that influenza infection increases plaque area, necrotic core size, and inflammatory cell infiltration, elevates chemokine protein levels in lung and aorta, induces Th17 responses, reduces anti-inflammatory macrophages, and directly infects vascular endothelial cells (11, 12, 14). However, the transcriptional programs underlying these cellular changes and the molecular basis for the persistence of cardiovascular risk beyond apparent clinical recovery remain undefined. Here we report comprehensive bulk RNA-seq analysis of aortic and lung tissues from HFD-fed *Apoe*−/− mice infected with influenza A/PR/8/34 versus PBS-treated controls at day 15 post-infection, the recovery phase at which cardiovascular risk remains elevated despite resolution of acute viral replication, aimed at identifying the molecular mechanisms by which pulmonary influenza infection drives atherosclerotic plaque destabilization

## 2. Materials and Methods

### 2.1 Animal Model and Experimental Design

*Apoe*−/− mice (10 weeks old, C57BL/6 background; The Jackson Laboratory) were maintained under specific pathogen-free conditions and fed a high-fat diet (HFD; 42% kcal fat, Teklad) for 8 weeks. Mice were then infected with influenza A/PR/8/34 (H1N1; 100 PFU) or PBS via oropharyngeal aspiration and euthanized on day 15 post-infection (Figure 1A). All procedures were approved by the University of Pittsburgh Institutional Animal Care and Use Committee.

**Figure 1.**
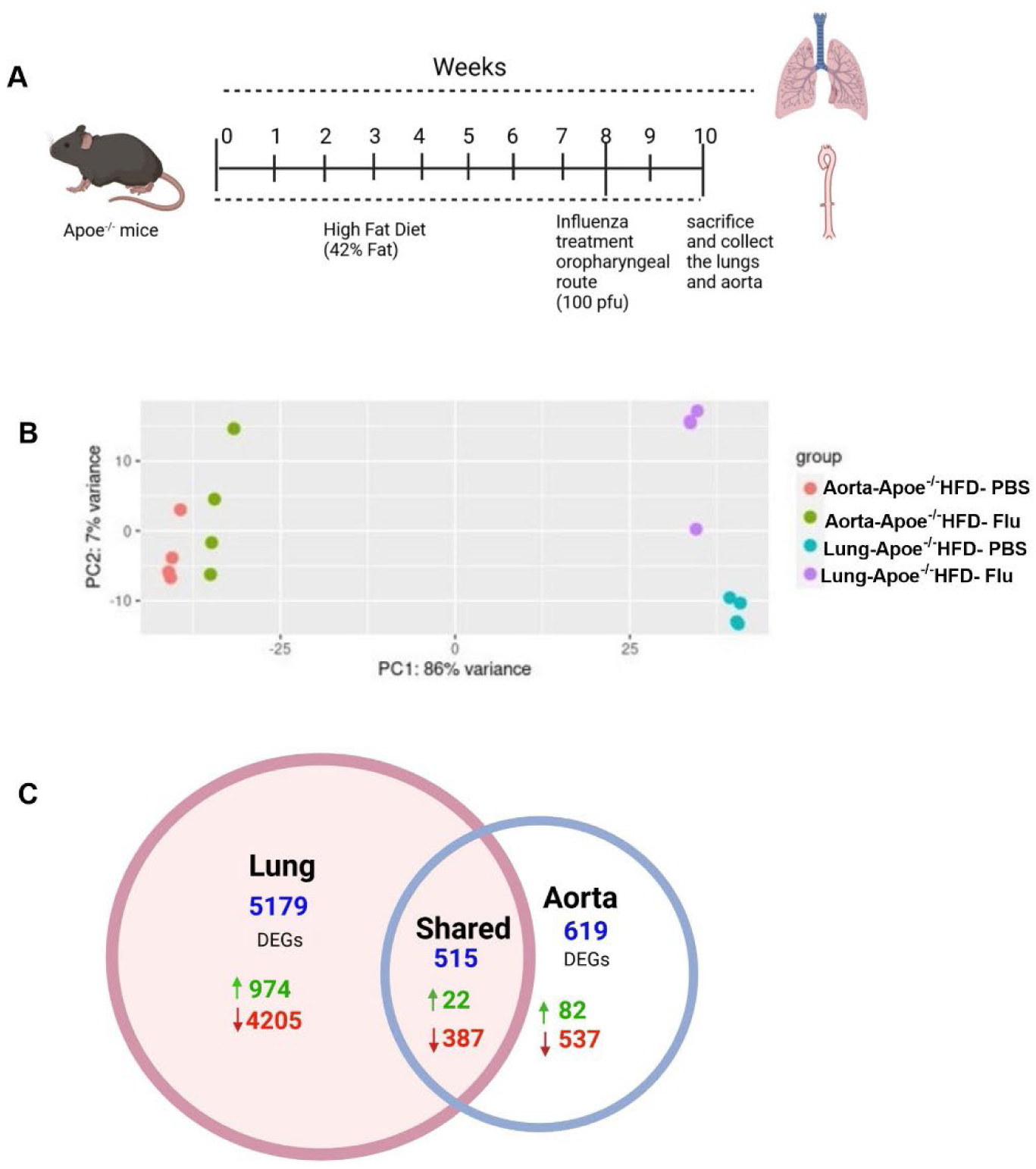
Genome-wide transcriptional response to influenza infection in *Apoe*−/− atherosclerotic mice. (A) Experimental design. *Apoe*−/− mice were fed a high-fat diet for 8 weeks and infected intranasally with influenza A/PR/8/34 (100 PFU) or PBS and sacrificed on day 15 post-infection (n=4 per group). Aorta and lung tissues were harvested for bulk RNA sequencing. (B) Principal component analysis of variance-stabilized RNA-seq counts. PC1 (86% variance) separates aortic from pulmonary samples, confirming tissue identity as the dominant source of transcriptional variance. PC2 (7% variance) captures the influenza treatment effect. Each dot represents one biological replicate. (C) Venn diagram showing overlap of differentially expressed genes across four gene lists: lung-upregulated, lung-downregulated, aorta-upregulated, and aorta-downregulated. HFD, high-fat diet; PBS, phosphate-buffered saline; PFU, plaque-forming unit.

### 2.2 Tissue Harvesting and RNA Extraction

Mice were perfused with 20 mL ice-cold PBS through the heart. The middle lung lobe and aorta (root to iliac bifurcation) were snap-frozen in liquid nitrogen and homogenized (14). RNA was isolated using the Absolutely RNA Miniprep Kit (Agilent Technologies).

### 2.3 RNA Sequencing and Differential Expression Analysis

RNA sequencing was performed by Genewiz (Azenta Life Sciences). RNA integrity and concentration were assessed by Agilent TapeStation and Qubit fluorometric assay. Libraries were prepared using a standard poly-A selection protocol, quantified by qPCR, and sequenced on an Illumina HiSeq platform in a 2×150 bp paired-end configuration with single indexing, generating approximately 350 million raw paired-end reads per lane. Raw sequencing data were provided in FASTQ format, and all downstream bioinformatics analyses were conducted by the Bioinformatics Core in the Department of Pediatrics using R version 4.3. Raw read quality was assessed using FastQC. Adapter sequences and low-quality bases were removed before alignment, and the quality of the trimmed reads was re-evaluated to confirm suitability for downstream analysis. Reads were aligned to the mouse reference genome (GRCm38/mm10), followed by gene-level count quantification. Principal component analysis was performed using variance-stabilized count data to evaluate sample quality and separation between experimental groups. Differential gene expression between influenza-infected and PBS-treated *Apoe*−/− mice maintained on a high-fat diet was assessed using DESeq2, with multiple-testing correction performed using the Benjamini-Hochberg method. Genes with an adjusted p-value < 0.05 and a log₂ fold change of at least 1.5 were classified as significantly upregulated, whereas genes with an adjusted p-value < 0.05 and a log₂ fold change below 0 were classified as significantly downregulated. Effect sizes are reported as log₂ fold changes (log₂FC). Cross-tissue overlap was assessed by calculating set intersections among four differential expression gene lists derived from the comparison of influenza-infected high-fat diet-fed *Apoe*−/− mice versus PBS-treated controls, comprising lung-upregulated, lung-downregulated, aorta-upregulated, and aorta-downregulated genes. Complete lists of differentially expressed genes for the lung and aorta, including log₂ fold change and adjusted p-values for all genes, are provided (Supplementary Data File).

### 2.4 Pathway Categorization and Quantitative RT-PCR Validation

Differentially expressed genes were assigned to biological pathway categories based on gene family and published function (Supplementary Table 1), and heatmaps were generated using log₂ fold change values. Gene expression was validated by quantitative reverse transcription PCR (qRT-PCR) using TaqMan probes and primers (Thermo Fisher Scientific, Waltham, MA, USA) (Supplementary Table 2). Fold changes in mRNA expression were calculated using the ΔΔCt method and normalized to the endogenous housekeeping gene hypoxanthine-guanine phosphoribosyltransferase (*Hprt*).

### 2.5 Bone Marrow-Derived Macrophage Generation and Influenza Treatment

Bone marrow-derived macrophages (BMDMs) were generated from *Apoe*−/− mice. Bone marrow was flushed from femora and tibiae under sterile conditions and cultured in DMEM supplemented with 10% fetal bovine serum, penicillin-streptomycin, and 20 ng/mL GM-CSF at 37°C in 5% CO₂ for seven days (14). Adherent BMDMs were replated for 24 hours and then treated with PBS or influenza A/PR/8/34 at MOI 1 for 24 hours, generating two experimental groups: *Apoe*−/− + PBS and *Apoe*−/− + influenza. Cell lysates were collected for gene expression analysis by qRT-PCR as described before (14).

### 2.6 Statistical Analysis

Data from qRT-PCR and BMDM experiments are presented as mean ± SEM. Statistical significance was determined using an unpaired Student’s t-test. Analyses were performed using GraphPad Prism. For the RNA-seq analysis, statistical thresholds are described in section 2.3. A p-value of less than 0.05 was considered statistically significant for all analyses.

## 3. Results

### 3.1 Overview of the Transcriptional Response to Influenza Infection

To characterize the transcriptional impact of influenza on atherosclerotic tissues, bulk RNA sequencing was performed on aorta and lung at day 15 post-infection. Principal component analysis of variance-stabilized RNA-seq counts is shown in Figure 1B. PC1 explained 86% of total variance and completely separated aortic from pulmonary samples, confirming that tissue identity is the dominant source of transcriptional variance. PC2 (7% variance) captured the influenza treatment effect, with influenza-infected lung samples separating clearly from PBS-treated lung controls along this axis. No outlier samples were identified, validating the quality and reproducibility of all biological replicates.

Influenza infection in HFD-fed *Apoe*−/− mice induced a combined total of 6,828 DEGs across the aorta and lung at day 15 post-infection, the recovery-phase time point at which acute viral replication has largely resolved but transcriptional consequences of infection persist in both the pulmonary and vascular compartments. Cross-tissue comparison (Figure 1C) identified 5,179 DEGs unique to the lung (974 upregulated, 4,205 downregulated), 619 DEGs unique to the aorta (82 upregulated, 537 downregulated), and 515 DEGs shared between both tissues. The lung exhibited a markedly larger transcriptional response with 5,694 total DEGs compared with 1,134 in the aorta, representing a five-fold greater perturbation. The shared gene set reflects systemic suppression of homeostatic gene programs common to both tissues, while the tissue-specific DEGs point to compartment-specific inflammatory and regulatory responses to infection.

### 3.2 Lung Transcriptome: Interferon Signaling and Anti-Viral Immune Response

The lung transcriptome showed direct evidence of persistent interferon-stimulated gene (ISG) expression and innate antiviral signaling at the recovery phase of influenza infection (Figure 2A). *Zbp1* (+2.02) was significantly upregulated in response to influenza infection. ZBP1 specifically recognizes influenza viral RNA generated during replication and triggers innate immune signaling including necroptosis and NLRP3 inflammasome activation (15). *Rsad2*/Viperin (+1.62), a broad-spectrum antiviral ISG that inhibits influenza replication (16), was also significantly induced. The IFN-inducible p200 family members *Ifi204* (+1.56) and *Ifi205* (+1.57), which sense cytosolic nucleic acids released during viral infection, were both upregulated. *Apol11b* (+3.04), an IFN-inducible apolipoprotein L family member (17), was strongly induced in the lung alongside *Tnfrsf9*/4-1BB (+2.71), indicating viral antigen-activated T and NK cells (18). These data suggest that innate antiviral and IFN signaling programs persist into the recovery phase, continuing to shape the pulmonary immune environment. IFN signaling was confirmed by qRT-PCR, with both *Zbp1* (Figure 2B) and *Apol11b* (Figure 2C) showing significant upregulation consistent with the RNA-seq findings.

**Figure 2.**
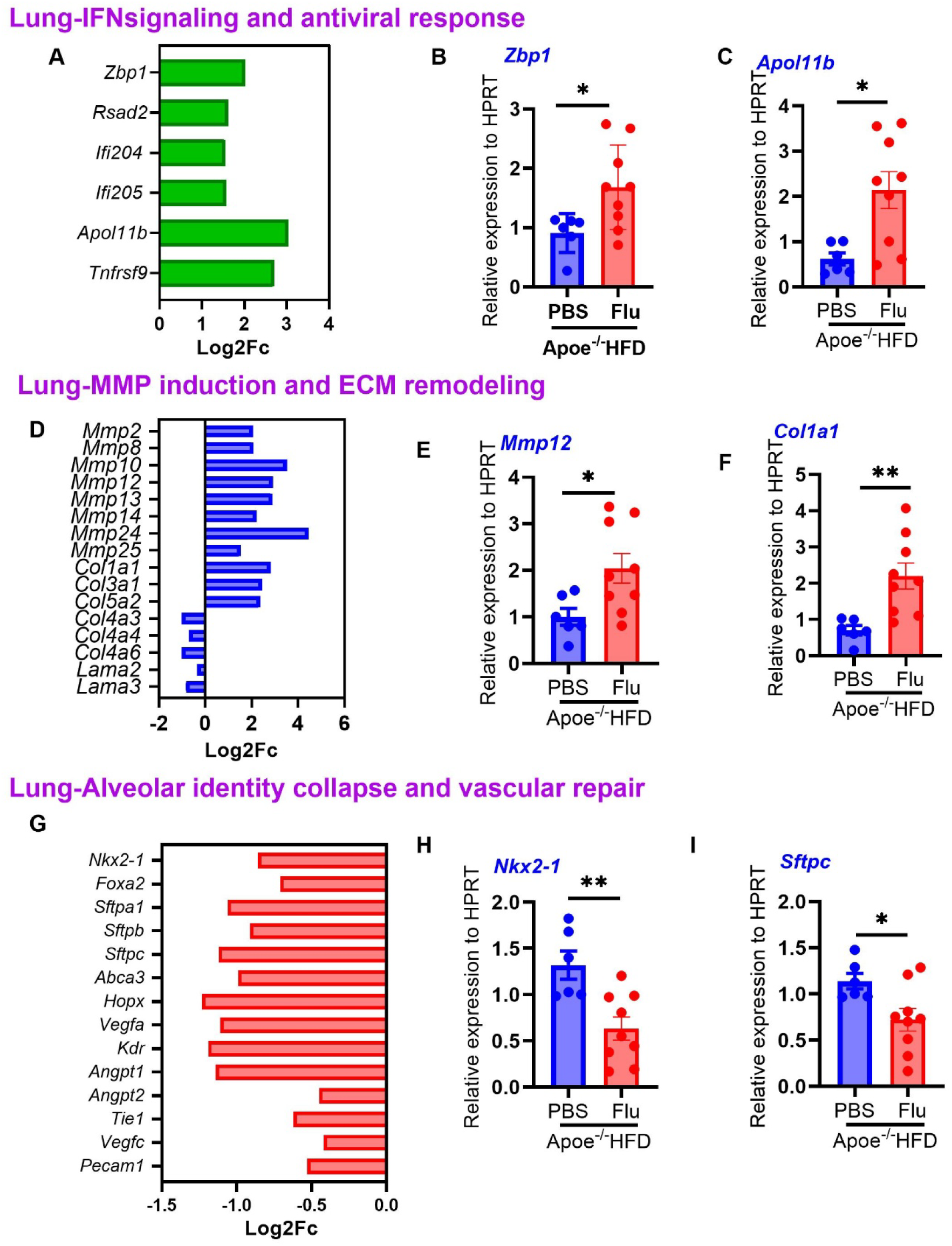
Influenza infection drives persistent IFN signaling, ECM remodeling, and alveolar identity collapse in the lung. (A) Heatmap showing log₂ fold change of interferon-stimulated and antiviral genes in the lung of influenza A/PR/8/34-infected versus PBS-treated high-fat diet-fed *Apoe*−/− mice at day 15 post-infection (adjusted p < 0.05). (B, C) Relative expression of *Zbp1* (B) and *Apol11b* (C) from lung tissue by qRT-PCR (n=6–9). (D) Heatmap of MMP/ECM remodeling genes. (E, F) Relative expression of *Mmp12* (E) and *Col1a1* (F) from lung tissue by qRT-PCR (n=6–9). (G) Heatmap of alveolar identity genes. (H, I) Relative expression of *Nkx2-1* (H) and *Sftpc* (I) from lung tissue by qRT-PCR (n=6–9). Data are presented as mean ± SEM. *p < 0.05 by unpaired two-tailed t-test. IFN, interferon; MMP, matrix metalloproteinase; ECM, extracellular matrix.

### 3.3 Lung Transcriptome: MMP Induction and ECM Remodeling

Influenza infection induced broad proteolytic remodeling of the pulmonary extracellular matrix. Eight matrix metalloproteinase genes were upregulated in the lung (Figure 2D) including *Mmp2* (+2.07), *Mmp8* (+2.08), *Mmp10* (+3.54), *Mmp12* (+2.92), *Mmp13* (+2.90), *Mmp14* (+2.21),

*Mmp24* (+4.46), and *Mmp25* (+1.54). This MMP induction is consistent with influenza-driven macrophage and neutrophil activation, both of which are major sources of matrix metalloproteinases in the infected lung (19, 20). Collagen and laminin genes showed bidirectional regulation, with upregulation of *Col1a1* (+2.82), *Col3a1* (+2.45), and *Col5a2* (+2.37) indicating fibrotic remodeling (19, 20), while downregulation of *Col4a3* (−1.01), *Col4a4* (−0.70), *Col4a6* (−1.01), *Lama2* (−0.35), and *Lama3* (−0.83) indicated basement membrane barrier disruption (19–21). This divergent ECM program reflects simultaneous fibrotic deposition and basement membrane disruption in the influenza-infected lung, consistent with the established role of macrophage and neutrophil-derived MMPs in pulmonary remodeling following viral infection (19, 20). MMP/ECM remodeling was confirmed by qRT-PCR, with *Mmp12* (Figure 2E) and *Col1a1* (Figure 2F) both showing significant upregulation consistent with the RNA-seq findings.

### 3.4 Lung Transcriptome: Alveolar Identity Collapse and Vascular Failure

Influenza infection imposed a coordinated suppression of alveolar epithelial identity and pulmonary vascular repair programs in the lung. Seven lung-identity genes were downregulated (Figure 2G) including the master alveolar transcription factor *Nkx2-1* (−0.86), *Foxa2* (−0.71), three surfactant protein genes *S*ftpa1 (−1.06), *Sftpb* (−0.91), and *Sftpc* (−1.12), *Abca3* (−0.99), and *Hopx* (−1.23) (22, 23). This coordinated downregulation of the alveolar type I and type II cell identity program indicates influenza-driven loss of alveolar epithelial integrity (24, 25). Seven angiogenesis genes were simultaneously downregulated including *Vegfa* (−1.11), *Kdr* (−1.19), *Angpt1* (−1.14), *Angpt2* (−0.45), *Tie1* (−0.62), *Vegfc* (−0.42), and *Pecam1* (−0.53), indicating collapse of the vascular repair program that would normally restore pulmonary vascular integrity following epithelial injury (26). These data suggest that influenza infection imposes a sustained state of alveolar structural compromise and vascular repair suppression at the recovery phase, providing a transcriptional basis for the persistence of pulmonary dysfunction beyond the acute phase. Alveolar identity collapse was confirmed by qRT-PCR, with *Nkx2-1* (Figure 2H) and *Sftpc* (Figure 2I) showing significant downregulation consistent with the RNA-seq findings.

### 3.5 Lung Transcriptome: Immune Cell Response

Influenza infection activated multiple innate and adaptive immune cell populations in the lung, establishing a systemic inflammatory program with direct relevance to atherosclerotic plaque progression. Macrophage and monocyte infiltration was reflected by upregulation of the pan-macrophage markers *Adgre1*/F4/80 (+2.26) and *Cd68* (+2.10)(27) (Figures 3A). *Trem2* (+3.97), a marker of lipid-associated macrophages, was among the most strongly induced macrophage genes, alongside the scavenger receptor *Msr1* (+3.34) indicating foam cell activity (28). Fc-γ receptors *Fcgr1* (+2.60), *Fcgr3* (+1.80), and *Fcgr2b* (+1.71) were all upregulated, indicating IgG immune complex-driven macrophage activation (29). *Nos2* (+1.60) and *Hmox1* (+2.31) reflect M1 macrophage activation, while *Arg1* (+4.71) and *Slc11a1* (+1.83) indicate concurrent M2-like and antimicrobial activity. The complete monocyte-mobilization program was present: *Ccl2* (+3.87), *Ccl7* (+4.12), *Ccl8* (+4.44), and *Ccl12* (+3.99), the four major murine CCR2 ligands. Ccl2 and Ccl7 are the dominant drivers of Ly6C-high monocyte egress from bone marrow (30–32).The pro-inflammatory cytokines *Tnf* (+2.24) and *Il6* (+1.72) were upregulated, reflecting macrophage and innate immune cell activation alongside the broader inflammatory response. Macrophage activation was confirmed by qRT-PCR, with *Il6* (Figure 3F), showing significant concordant differential expression.

**Figure 3.**
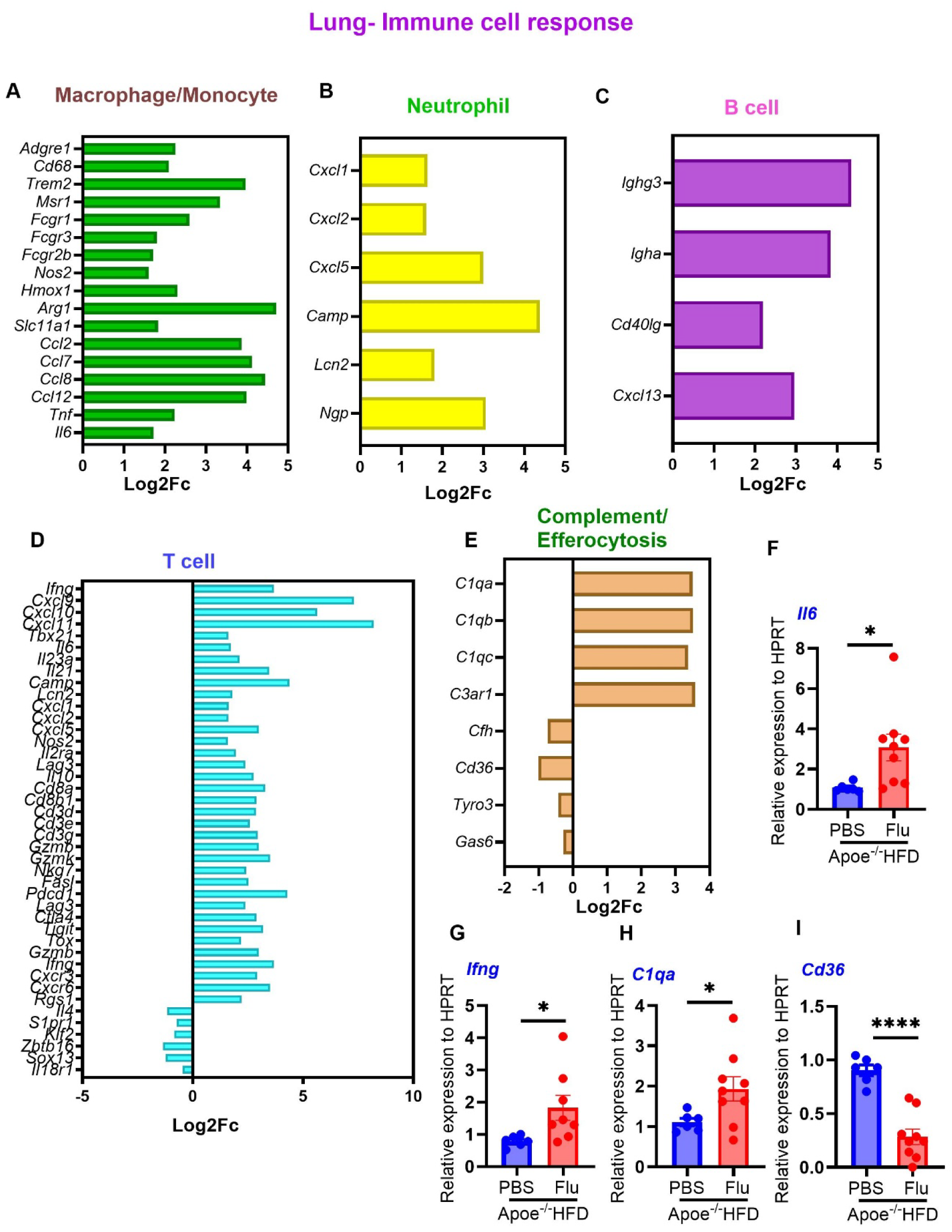
Influenza infection activates broad innate and adaptive immune cell programs in the lung. (A-E) Heatmap showing log₂ fold change of immune cell response genes organized by cell type, including macrophage/monocyte (A), neutrophil (B), and B cell (C) and T cell (D), and complement/efferocytosis genes (E) in the lung of influenza A/PR/8/34-infected versus PBS-treated high-fat diet-fed *Apoe*−/− mice at day 15 post-infection (adjusted p < 0.05). Relative expression of *Il6* (F), *Ifng* (G), *C1qa* (H) and *Cd36* (I) from lung tissue by qRT-PCR (n=6–9). Data are presented as mean ± SEM. *p < 0.05 by unpaired two-tailed t-test.

Neutrophil recruitment signals were strongly activated. *Cxcl1* (+1.64), *Cxcl2* (+1.61), and *Cxcl5* (+3.00) are CXCR2 ligands that recruit neutrophils to sites of inflammation and are direct IL-17 transcriptional targets (33) (Figure 3B). *Camp* (cathelicidin, +4.38) and *Lcn2* (+1.81) are neutrophil antimicrobial effectors, and *Ngp* (+3.06) is a specific granule marker (34).

B-cell activity was reflected by upregulation of the class-switched immunoglobulin heavy chains *Ighg3* (+4.35) and *Igha* (+3.85), indicating IgG3-and IgA-producing B-cell responses within the lung (35) (Figure 3C). *Cd40lg* (+2.20) supports T-follicular helper cell activity driving B-cell class-switch recombination, and *Cxcl13* (+2.96) indicates B-cell follicle organization consistent with ectopic lymphoid structure formation (36).

T-cell responses were broadly activated across multiple effector subsets, with concurrent exhaustion checkpoint induction consistent with the day 15 post-infection time point (Figure 3D). The Th1 program was dominant, with *Ifng* (+3.68) driving *Cxcl9* (+7.32), *Cxcl10* (+5.65), and *Cxcl11* (+8.21) in a self-amplifying CXCR3 feed-forward loop and the Th1 master transcription factor *Tbx21*/T-bet (+1.61) confirming a polarized Th1 effector response (37, 38). The Th2 program was actively suppressed, with *Il4* (−1.16) significantly downregulated, suggesting influenza-driven suppression of type 2 immunity (38). The Th17 program was comprehensively activated, with *Il6* (+1.72) (Figure 3A) and *Il23a* (+2.12) driving differentiation, *Il21* (+3.48) providing autocrine amplification, and downstream effectors *Camp* (+4.38), *Lcn2* (+1.81), *Cxcl1* (+1.64), *Cxcl2* (+1.61), *Cxcl5* (+3.00), and *Nos2* (+1.60) confirming active IL-17 signaling, consistent with elevated CD4+RORγt+ cells and serum IL-17 previously reported in this model (14). A partial Treg signature was present through *Il2ra* (+1.96) and *Il10* (+2.76), with Tr1-like counter-regulation reflected by *Lag3* and *Il10* co-expression rather than canonical *Foxp3*-dependent suppression (39). A substantial CD8^+^ T-cell infiltrate was confirmed by *Cd8a* (+3.28), *Cd8b1* (+2.89), and T-cell receptor complex components *Cd3d* (+2.87), *Cd3e* (+2.59), and *Cd3g* (+2.94). The cytotoxic effector program was activated through *Gzmb* (+3.00), *Gzmk* (+3.51), *Nkg7* (+2.42), and *Fasl* (+2.52), while exhaustion checkpoints *Pdcd1*/PD-1 (+4.29), *Lag3* (+2.38), *Ctla4* (+2.90), *Tigit* (+3.18), and *Tox* (+2.20) were simultaneously induced, defining a transitional effector-to-exhaustion state at day 15 post-infection (40). The memory T-cell compartment showed a three-way split. Central memory recirculation was being lost through *S1pr1* (−0.73) and *Klf2* (−0.85), effector memory was strongly active through *Gzmb*, *Ifng*, and *Cxcr3*, and tissue-resident memory was emerging through *Cxcr6* (+3.51) and *Rgs1* (+2.21), consistent with TRM seeding (41, 42). Innate-like T-cell populations were broadly suppressed. *Zbtb16*/PLZF (−1.34), the shared master transcription factor of NKT, iNKT, and MAIT cells, was downregulated alongside the γδ T-cell master transcription factor *Sox13* (−1.24) and *Il18r1* (−0.46), indicating a shift from innate-like to adaptive T-cell dominance (43, 44). Together these findings define a pulmonary T-cell landscape characterized by dominant Th1 and Th17 effector activity, active CD8^+^ cytotoxic killing with emerging exhaustion, and tissue-resident memory seeding, reflecting a sustained adaptive immune response in the influenza-infected lung that persists into the recovery phase despite reduced viral burden. T-cell activation was confirmed by qRT-PCR, with *Ifng* (Figure 3G) showing significant upregulation consistent with the RNA-seq findings.

The complement system was strongly activated in the lung. *C1qa* (+3.50), *C1qb* (+3.52), and *C1qc* (+3.37) were among the most significantly upregulated genes alongside *C3ar1* (+3.57), while *Cfh* (−0.74), the principal alternative pathway inhibitor, was simultaneously downregulated, further amplifying complement activation (45) (Figure 3E). Efferocytosis was simultaneously impaired through downregulation of *Cd36* (−1.01), *Tyro3* (−0.43), and the bridging ligand *Gas6* (−0.28), allowing apoptotic cells to undergo secondary necrosis and contributing to tissue damage in the infected lung (46, 47). These data suggest that influenza infection drives sustained complement activation and efferocytosis failure in the lung, impairing the clearance of apoptotic cells and perpetuating the pulmonary inflammatory response. Complement activation and efferocytosis impairment were confirmed by qRT-PCR, with *C1qa* (Figure 3H) showing significant upregulation and *Cd36* (Figure 3I) showing significant downregulation consistent with the RNA-seq findings.

### 3.6 Lung Transcriptome: Acute Phase Response and Oxidative Stress

Influenza infection induced a marked acute phase response in the lung (Figure 4A). *Saa3* (+6.36) and *Saa1* (+3.75), among the most sensitive acute phase reactants, remodel HDL toward a pro-inflammatory phenotype, promote foam cell formation, and induce endothelial adhesion molecule expression (48–50). *A2m* (+5.32), a broad-spectrum protease inhibitor, reflects the magnitude of the systemic cytokine-driven response (51). *Ptx3* (+3.15), a long pentraxin acute phase protein, functions in complement activation and pathogen recognition at sites of local inflammation (52, 53). *Lcn2* (+1.81) contributes both as an IL-17 downstream effector and an acute phase protein (54).

**Figure 4.**
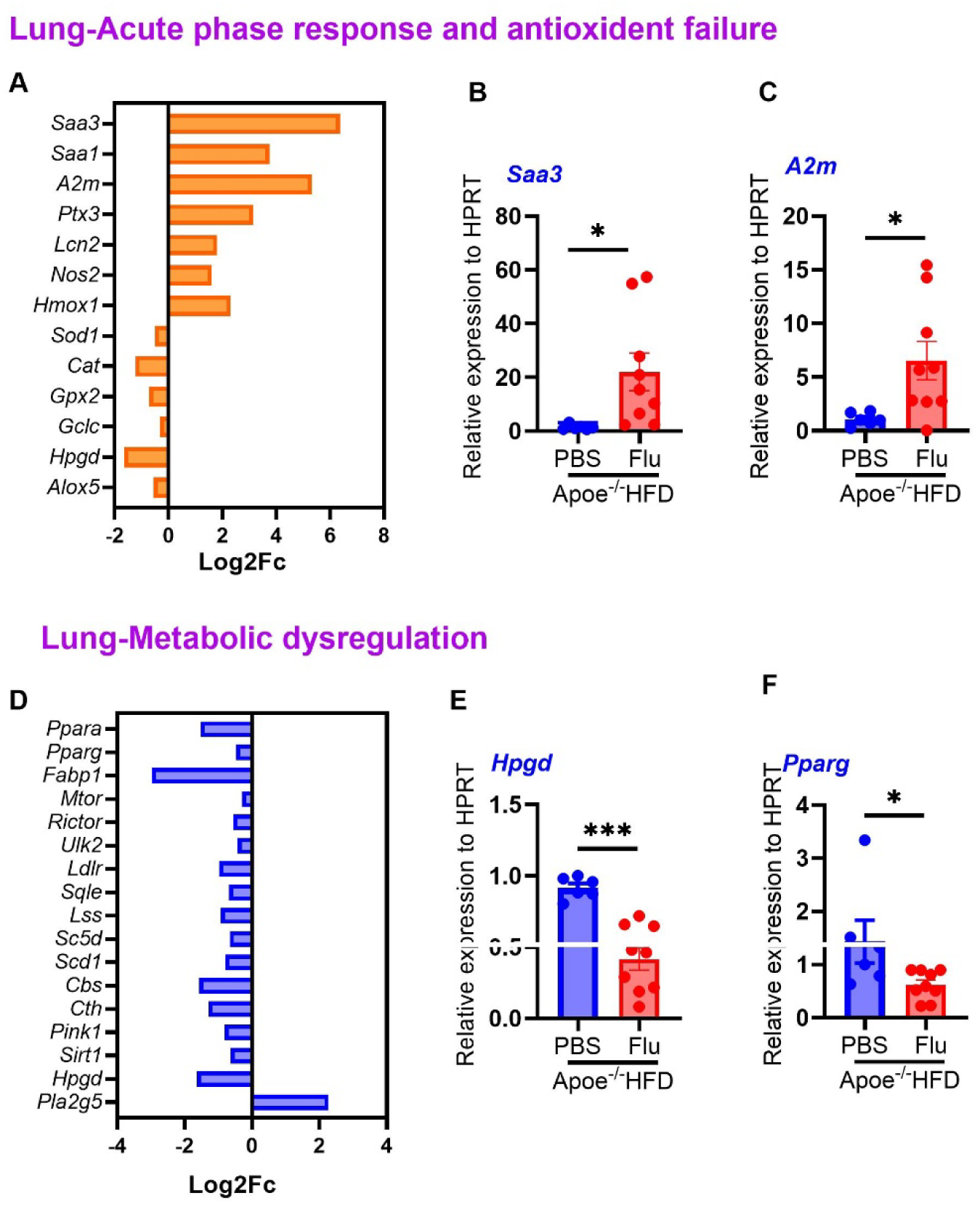
Influenza infection induces sustained acute phase response and metabolic dysregulation in the lung. (A) Heatmap showing log₂ fold change of acute phase and oxidative stress genes in the lung of influenza A/PR/8/34-infected versus PBS-treated high-fat diet-fed *Apoe*−/− mice at day 15 post-infection (adjusted p < 0.05). (B, C) Relative expression of *Saa3* (B) and *A2m* (C) from lung tissue by qRT-PCR (n=6–9), reflecting the acute phase response. (D) Heatmap of metabolic dysregulation genes. (E, F) Relative expression of *Pparg* (E) and *Hpgd* (F) from lung tissue by qRT-PCR (n=6–9), reflecting metabolic dysregulation. Data are presented as mean ± SEM. *p < 0.05 by unpaired two-tailed t-test.

The oxidative stress landscape showed a divergent pattern of simultaneous ROS production and antioxidant defense failure. *Nos2*/iNOS (+1.60) and *Hmox1* (+2.31) were upregulated, reflecting M1 macrophage activation and cytoprotective stress response (55), while antioxidant enzymes *Sod1* (−0.50), *Cat* (−1.23), *Gpx2* (−0.71), and the glutathione synthesis enzyme *Gclc* (−0.31) were concurrently suppressed, indicating uncompensated oxidative stress that would amplify inflammatory tissue damage beyond the acute phase (55).

The inflammatory resolution program was also compromised. *Hpgd* (−1.64), the principal prostaglandin-inactivating enzyme, was downregulated, allowing pro-inflammatory prostaglandins to accumulate and prolong inflammatory signaling (56). Alox5 (−0.56) downregulation indicates suppression of the 5-lipoxygenase eicosanoid pathway, which through reduced lipoxin generation may impair inflammatory resolution (57). These data suggest that the pulmonary acute phase response, oxidative stress, and failure of inflammatory resolution collectively create a self-sustaining pro-inflammatory state in the influenza-infected lung. Acute phase findings were confirmed by qRT-PCR, with *Saa3* (Figure 4B) and *A2m* (Figure 4C) showing significant upregulation consistent with the RNA-seq findings.

### 3.7 Lung Transcriptome: Metabolic Dysregulation

Influenza infection imposed broad suppression of metabolic pathways in the lung (Figure 4D), impairing the metabolic flexibility required for inflammatory resolution. Fatty acid metabolism was impaired through downregulation of *Ppara* (−1.53) and *Pparg* (−0.47), mechanistically linked to loss of M2 macrophage polarization capacity (58), and the lipid transport protein *Fabp1* (−2.97). Fifteen mitochondrial OXPHOS genes and 13 glycolytic enzymes were downregulated, reflecting widespread bioenergetic failure. *Mtor* (−0.30), *Rictor* (−0.56), and *Ulk2* (−0.42) suppression indicates attenuation of mTORC1/2-dependent metabolic signaling (59), while *Ulk2* suppression simultaneously impairs autophagy initiation, compounding mitophagic failure (60).

Cholesterol uptake was impaired through *Ldlr* (−0.97) downregulation, while biosynthesis was broadly suppressed through *Sqle* (−0.68), *Lss* (−0.92), and *Sc5d* (−0.66). *Scd1* (−0.79) suppression increased the saturated to monounsaturated fatty acid ratio, directly promoting ER stress and NLRP3 inflammasome priming (61). The transsulfuration pathway was impaired through *Cbs* (−1.57) and *Cth* (−1.29). *Cbs* suppression causes homocysteine accumulation, promoting endothelial dysfunction and foam cell formation (62), while *Cth* suppression depletes cysteine availability for glutathione synthesis, impairing antioxidant defense and amplifying oxidative stress. *Pink1* (−0.81) suppression impairs mitophagy (63), allowing damaged mitochondria to release DAMPs that sustain innate immune activation. *Sirt1* (−0.64) suppression removes the transcriptional brake on NF-κB inflammatory signaling (64). *Pla2g5* (+2.27) upregulation increases arachidonic acid liberation, while *Hpgd* (−1.64) suppression impairs prostaglandin degradation, creating a two-hit eicosanoid imbalance that favors sustained pro-inflammatory signaling (65). These data suggest that influenza infection imposes a state of metabolic collapse in the lung, impairing the bioenergetic and biosynthetic flexibility required for inflammatory resolution and tissue repair. Metabolic dysregulation was confirmed by qRT-PCR, with *Pparg* (Figure 4E) and *Hpgd* (Figure 4F) showing significant downregulation consistent with the RNA-seq findings.

### 3.8 Aortic Transcriptome: Interferon Signaling and Anti-Viral Immune Response

In addition to the pulmonary IFN response, direct transcriptional evidence of IFN signaling was detected in the aorta (Figure 5A), consistent with influenza M protein expression and Color-flu Venus-positive endothelial cells confirmed in this model (14). *Slfn4* (+1.89), a Type I IFN-stimulated gene with restricted expression in monocytes and macrophages, indicates IFN-α/β signaling within the macrophage-rich plaque (66). *Apol11b* (+5.10 aorta versus +3.04 lung) is more strongly induced in the aorta than the lung, indicating a local rather than purely systemic antiviral response at the vessel wall (67). *Tnfrsf9*/4-1BB (+1.88) suggests the presence of viral antigen-activated T or NK cells, and *Gzmb* (+2.85) indicates active cytotoxic killing within the plaque (68, 69). These data suggest that influenza infection directly engages the aortic wall as an active site of antiviral immune signaling, with local IFN-α/β induction and viral antigen-activated cytotoxic cells persisting during the recovery phase. Local IFN signaling was confirmed by qRT-PCR, with *Apol11b* showing significant upregulation consistent with the RNA-seq findings (Figure 5B).

**Figure 5.**
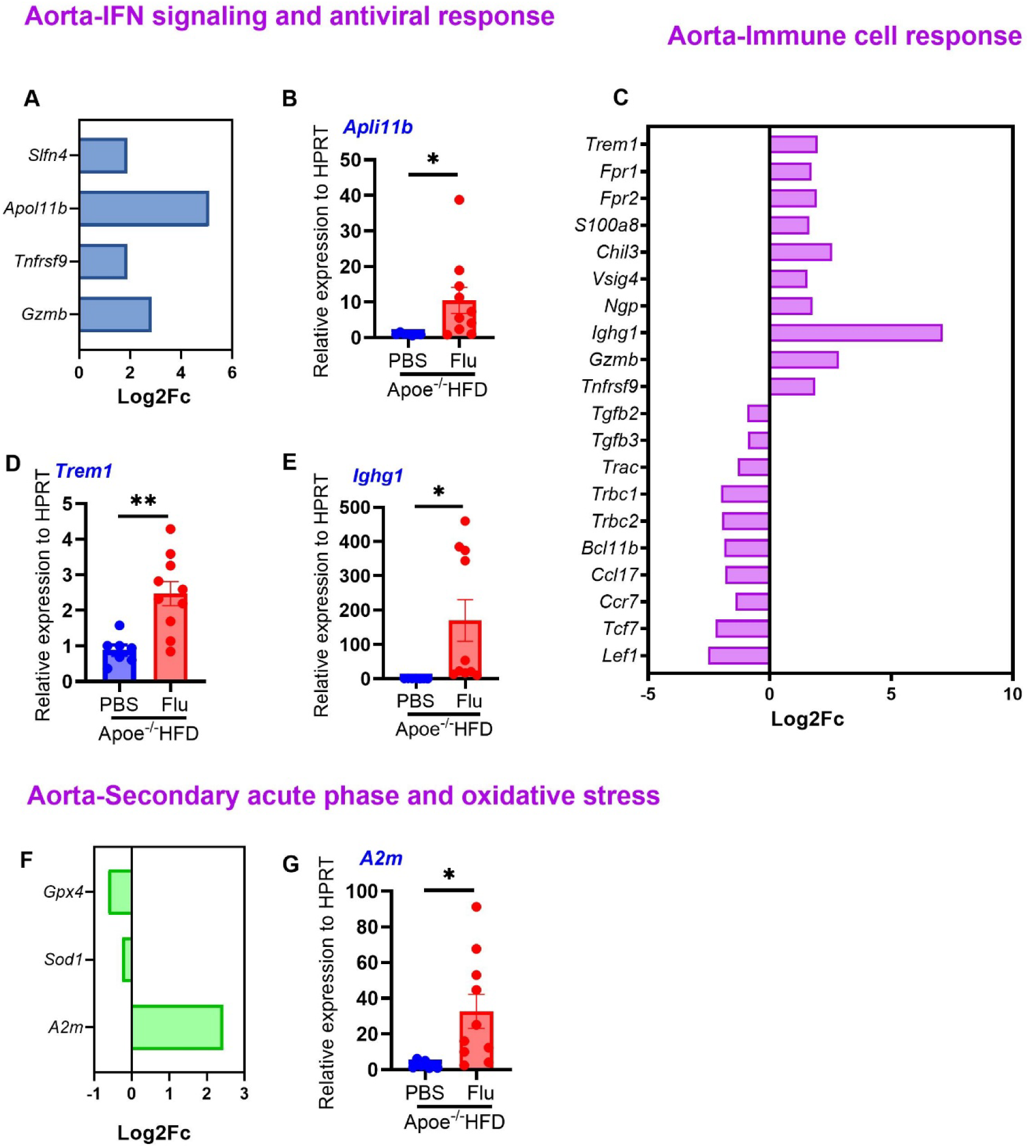
Influenza infection drives immune activation, and acute phase response gene expression in the aorta. (A) Heatmap showing log₂ fold change of IFN/antiviral genes in the aorta of influenza A/PR/8/34-infected versus PBS-treated high-fat diet-fed *Apoe*−/− mice at day 15 post-infection (adjusted p < 0.05). (B) Relative expression of *Apol11b* from aortic tissue. (C) Heatmap of aortic immune cell response genes. (D, E) Relative expression of *Trem1* (D) and *Ighg1* (E) from aortic tissue. (F) Heatmap of aortic acute phase and oxidative stress genes. (G) Relative expression of *A2m* from aortic tissue (n=7–10). Data are presented as mean ± SEM. *p < 0.05 by unpaired two-tailed t-test.

### 3.9 Aortic Transcriptome: Immune Cell Response

The aortic transcriptome revealed a broad immune cell response across innate and adaptive compartments, reflecting the systemic inflammatory response to influenza infection (Figure 5C). Macrophage and monocyte activation was evident in the aorta, consistent with the inflammatory cell infiltration documented in this model (14). *Trem1* (+1.97), a myeloid activation amplifier, was upregulated alongside the formyl peptide receptors *Fpr1* (+1.74) and *Fpr2* (+1.94), which respond to mitochondrial DAMPs released from necrotic plaque cells (70, 71). *S100a8* (+1.63), a potent DAMP and RAGE ligand, amplifies local TLR4 and NF-κB signaling, and together with *Trem1* and *Fpr1/Fpr2* suggests a feed-forward DAMP signaling loop amplifying NF-κB-driven vascular inflammation at the plaque microenvironment (71). *Chil3*/YM1 (+2.57) and *Vsig4* (+1.56) mark resident plaque macrophage populations (72, 73). Loss of *Tgfb2* (−0.91) and *Tgfb3* (−0.90) removes the TGF-β quiescence signal that normally promotes M2 macrophage polarization, consistent with the decreased CD206+F4/80+ anti-inflammatory macrophages documented in this model (14). Neutrophil presence within the aortic wall is reflected by *Ngp* (+1.76), a specific granule marker indicating neutrophil infiltration into the plaque microenvironment, consistent with the chemokine-driven neutrophil recruitment signals active in the lung (Figure 3A). Macrophage activation was confirmed by qRT-PCR, with *Trem1* showing significant upregulation consistent with the RNA-seq findings (Figure 5D).

B-cell humoral activity was strongly represented in the aorta (Figure 5C). Thirty-one immunoglobulin variable region genes were upregulated including multiple *Igkv* and *Ighv* family members, and *Ighg1* (+7.13) suggests class-switched, antigen-experienced B cells or plasma cells are present at the aortic wall (35). Humoral activation was confirmed by qRT-PCR, with *Ighg1* showing significant upregulation consistent with the RNA-seq findings (Figure 5E).

T-cell and lymphocyte responses in the aorta showed active effector engagement with concurrent remodeling of the regulatory landscape (Figure 5C). Cytotoxic immune activity was confirmed by *Gzmb* (+2.85) and *Tnfrsf9*/4-1BB (+1.88), consistent with the increased CD8+ and CD4+ T cells documented by flow cytometry (14). Although T-cell receptor components *Trac* (−1.30), *Trbc1* (−2.00), *Trbc2* (−1.97), and *Bcl11b* (−1.87) were downregulated, the strong induction of *Gzmb* (+2.85) suggests active cytotoxic T-cell engagement within the aortic wall (69). Th17 functional activity was evidenced by *S100a8* (+1.63), induced in aortic myeloid cells by IL-17A, consistent with elevated serum IL-17 and increased CD4+RORγt+ cells in this model (14). Treg recruitment was reduced through *Ccl17* (−1.83), the principal CCR4 ligand attracting regulatory T cells (74). Downregulation of the lymphatic egress receptor *Ccr7* (−1.40) alongside *Tcf7* (−2.22) and *Lef1* (−2.52) indicates that T cells recruited to the aortic wall are retained within the tissue and do not recirculate, consistent with establishment of a local effector T-cell population at the site of vascular inflammation. These data suggest that the aortic immune compartment has transitioned from homeostatic surveillance to active inflammatory engagement, with recruited macrophages, cytotoxic T cells, and class-switched B cells collectively sustaining plaque inflammation, while the concurrent loss of Treg recruitment and retention of effector T cells within the tissue creates an unregulated inflammatory microenvironment that favors progressive plaque destabilization.

### 3.10 Aortic Transcriptome: Acute Phase Response and Oxidative Stress

The aorta showed transcriptional evidence of a secondary acute phase response (Figure 5F). *A2m* (+2.44) was significantly upregulated, indicating that the systemic acute phase response originating in the lung has activated vascular acute phase gene expression in the aortic compartment (75). Antioxidant defenses were simultaneously compromised. *Sod1* (−0.26) is a frontline defense against superoxide-mediated vascular damage (76), while *Gpx4* (−0.62) is the principal enzyme protecting against ferroptosis (77), its suppression creating a permissive environment for lipid peroxidation and oxidative plaque destabilization (76, 78). These data suggest that the aortic plaque is simultaneously exposed to systemic acute phase signals from the lung and deprived of its antioxidant defenses, creating conditions for oxidative injury and ferroptotic cell death within the plaque. The secondary acute phase response was confirmed by qRT-PCR, with A2m showing significant upregulation consistent with the RNA-seq findings (Figure 5G).

### 3.11 Aortic Transcriptome: Metabolic Dysregulation

The aortic metabolic state was severely compromised through convergent failure of multiple energy pathways (Figure 6A). Twenty-four mitochondrial OXPHOS genes were downregulated, and glycolysis was simultaneously suppressed through *Hk2* (−1.34), *Aldoa* (−0.72), and *Pkm* (−0.46), representing complete bioenergetic failure in aortic plaque cells. *Acaca* (−2.32) indicated fatty acid synthesis blockade, removing the principal substrate for membrane lipid synthesis and macrophage function (79). *Scd1* (−1.79) suppression increased membrane saturated fatty acid content, directly promoting macrophage ER stress and NLRP3 inflammasome priming (80). *Ldlr* (−1.19) downregulation further reduced local LDL clearance in an already hypercholesterolemic environment, accelerating lipid accumulation and foam cell formation within the plaque (81). *Pink1* (−0.39) suppression impairs mitophagy, allowing damaged mitochondria to release mitochondrial DAMPs that sustain innate immune activation and NLRP3 inflammasome assembly within the plaque (82). *Bdh1* (−1.01) downregulation eliminated ketone body oxidation as the last remaining alternative fuel source, completing the bioenergetic collapse and removing the anti-inflammatory brake that β-hydroxybutyrate normally exerts on NLRP3 inflammasome activation (83). Together, these findings define an aortic plaque environment of convergent metabolic failure, with bioenergetic collapse, cholesterol dysregulation, mitophagic impairment, and NLRP3 inflammasome priming collectively creating conditions that favor necrotic core expansion and plaque instability. Aortic metabolic dysregulation was confirmed by qRT-PCR, with *Scd1* showing significant downregulation (Figure 6B).

**Figure 6.**
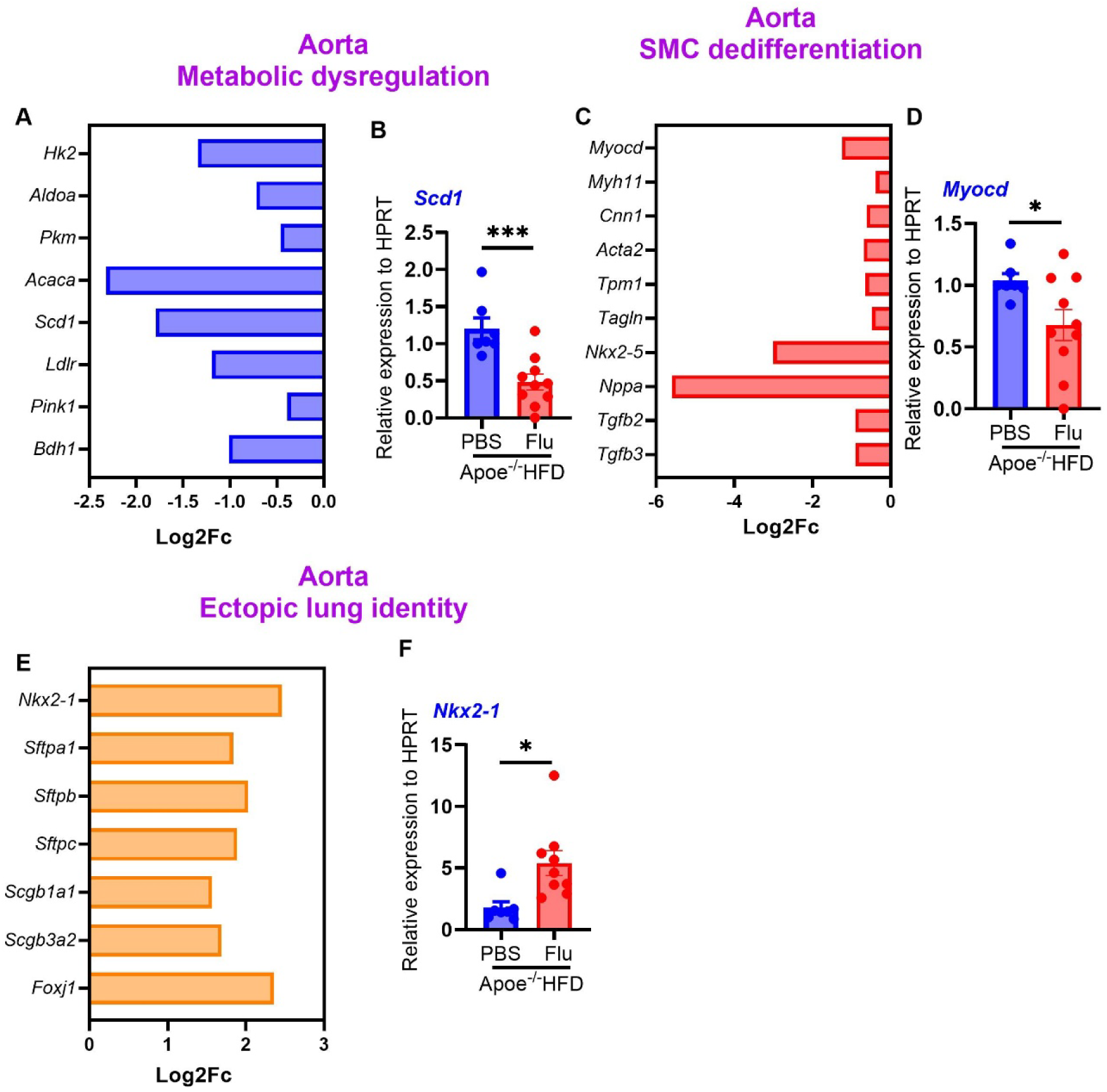
Influenza infection drives metabolic failure, SMC dedifferentiation, and ectopic lung-identity expression in the aorta. (A) Heatmap showing log₂ fold change of metabolic dysregulation genes in the aorta. (B) Relative expression of *Scd1* from aortic tissue. (C) Heatmap of SMC dedifferentiation genes. (D) Relative expression of *Myocd* from aortic tissue. (E) Heatmap of ectopic lung-identity genes upregulated in the aorta. (F) Relative expression of *Nkx2-1* from aortic tissue. All qRT-PCR data from aortic tissue (n=7–10). Data are presented as mean ± SEM. *p < 0.05 by unpaired two-tailed t-test. SMC, smooth muscle cell.

### 3.12 Aortic Transcriptome: Smooth Muscle Cell Dedifferentiation and Structural Compromise

Influenza infection drove coordinated suppression of the smooth muscle cell contractile identity program in the aorta. Nine SMC identity genes were downregulated (Figure 6C). The master SMC regulator *Myocd* (−1.25) was suppressed alongside contractile program genes *Myh11* (−0.38), *Cnn1* (−0.62), *Acta2* (−0.69), *Tpm1* (−0.65), and *Tagln* (−0.48), and the vascular identity genes *Nkx2-5* (−3.01) and *Nppa* (−5.59). Loss of the SMC quiescence-maintaining factors *Tgfb2* (−0.91) and *Tgfb3* (−0.90) removes the TGF-β brake on SMC phenotypic switching (84). These data suggest that the loss of the SMC contractile program transitions smooth muscle cells toward a synthetic, pro-inflammatory state, reducing fibrous cap integrity and increasing structural vulnerability to plaque rupture. SMC dedifferentiation was confirmed by qRT-PCR, with *Myocd* showing significant downregulation consistent with the RNA-seq findings (Figure 6D).

### 3.13 Ectopic Lung-Identity Genes in the Aorta

A striking and unexpected finding in the aortic transcriptome was the upregulation of seven lung-identity genes spanning three distinct lung cell types (Figure 6E). These included *Nkx2-1* (+2.46), the master transcription factor for lung epithelial identity, the alveolar type II surfactant genes Sftpa1 (+1.84), *Sftpb* (+2.03), and *Sftpc* (+1.89), the club cell markers *Scgb1a1* (+1.57) and *Scgb3a2* (+1.69), and the ciliated cell transcription factor *Foxj1* (+2.36) (85). These genes have no established functional role in the aorta and were simultaneously downregulated in the lung, representing a fully discordant, tissue-opposing regulatory pattern. These findings identify an ectopic lung-identity transcriptional signature in the aorta as a novel feature of influenza-exacerbated atherosclerosis, the mechanistic basis of which warrants direct experimental investigation. Ectopic lung-identity expression in the aorta was confirmed by qRT-PCR, with *Nkx2-1* showing significant upregulation consistent with the RNA-seq findings (Figure 6F).

### 3.14 Lung and Aortic Shared Transcriptome: Circadian Clock Suppression

Influenza infection induced broad suppression of the molecular circadian clock in the lung (Figure 7A). Seven circadian genes were downregulated including *Nr1d1*/REV-ERBα (−1.86), *Nr1d2*/REV-ERBβ (−0.60), *Per1* (−0.91), *Per3* (−0.56), *Cry2* (−1.05), *Arntl*/BMAL1 (−0.68), and *Clock* (−0.75) (86–88). The BMAL1/CLOCK transcriptional activators and the REV-ERB/PER/CRY repressive limb of the circadian feedback loop were collectively suppressed, representing coordinated suppression of the entire core clock mechanism in the lung (89). Disruption of *Nr1d1*/REV-ERBα removes the transcriptional brake that normally represses inflammatory gene expression during the rest phase (90), and *Arntl*/BMAL1 suppression amplifies NF-κB signaling and macrophage inflammatory polarization(91). Circadian clock suppression in the lung was confirmed by qRT-PCR, with *Nr1d1* showing significant downregulation consistent with the RNA-seq findings (Figure 7B).

**Figure 7.**
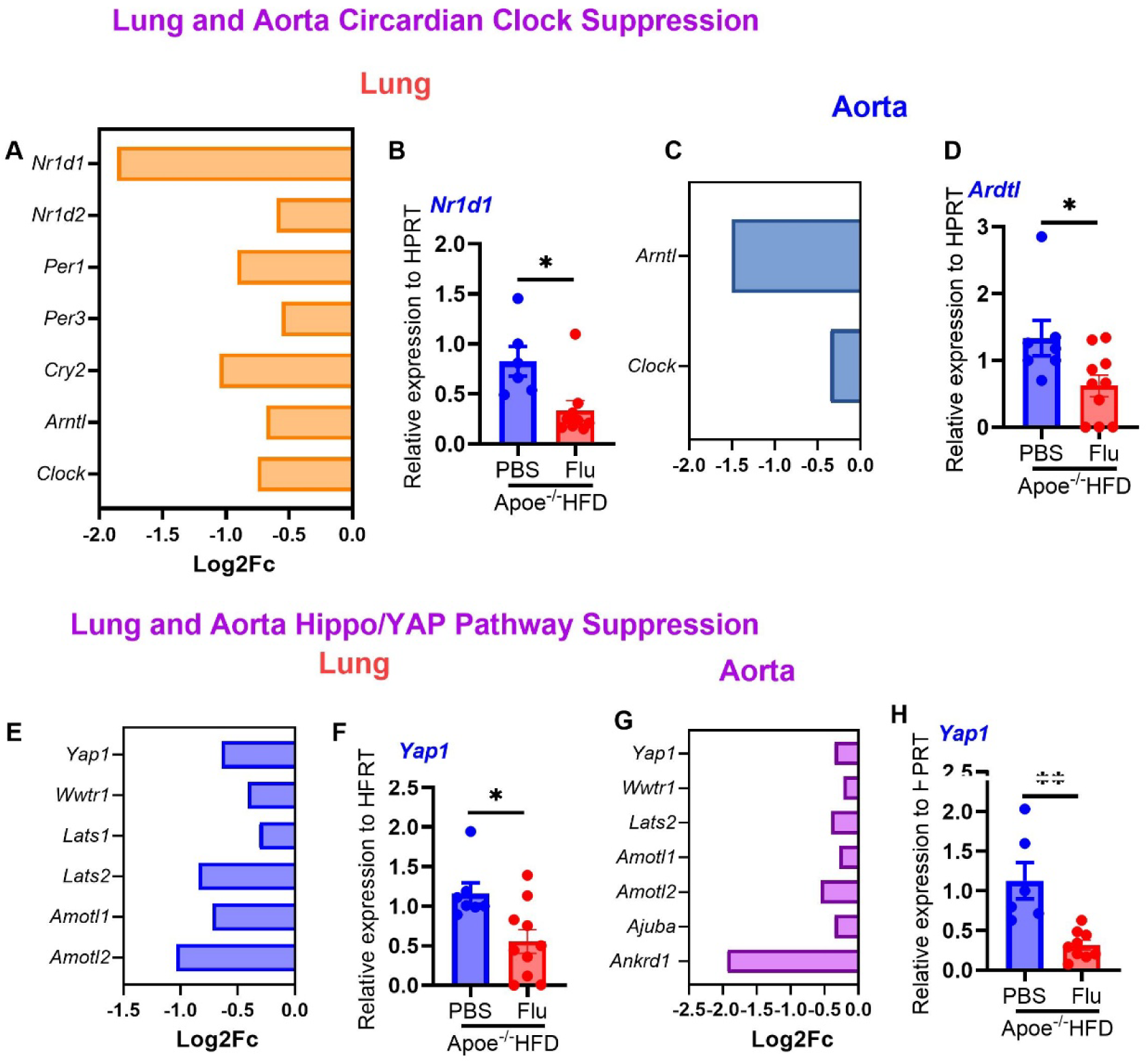
Influenza infection suppresses circadian clock and Hippo/YAP mechanosensing pathways in both lung and aorta. (A) Heatmap showing log₂ fold change of circadian clock genes in the lung of influenza A/PR/8/34-infected versus PBS-treated high-fat diet-fed *Apoe*−/− mice at day 15 post-infection (adjusted p < 0.05). (B) Relative expression of *Nr1d1* from lung tissue (n=6–9). (C) Heatmap showing log₂ fold change of circadian clock genes in the aorta (D) Relative expression of *Arntl* from aortic tissue by qRT-PCR (n=7–10). (E) Heatmap of Hippo/YAP pathway genes in lung. (F) Relative expression of *Yap1* from lung tissue (n=6-9). (G) Heatmap of Hippo/YAP pathway genes in the aorta. (H) Relative expression of *Yap1* from aortic tissue by qRT-PCR (n=7–10). Data are presented as mean ± SEM. *p < 0.05 by unpaired two-tailed t-test. YAP, yes-associated protein.

In the aorta, the same core clock components were suppressed, indicating that circadian disruption extends beyond the lung to the vascular compartment. *Arntl*/BMAL1 (−1.50) and *Clock* (−0.35) were both downregulated, and circadian regulation of endothelial shear stress responses, macrophage inflammatory capacity, and T-cell trafficking all depend on intact BMAL1/CLOCK function (90, 91) (Figure 7C). Their suppression in the aorta removes the temporal gating that normally restrains vascular inflammation. These data suggest that influenza infection imposes circadian clock suppression across both tissues simultaneously, removing the temporal restraint on vascular inflammation and amplifying the inflammatory programs identified in the aortic transcriptome. Circadian clock suppression in the aorta was confirmed by qRT-PCR, with *Arntl* showing significant downregulation consistent with the RNA-seq findings (Figure 7D).

### 3.15 Lung and Aortic Shared Transcriptome: Hippo/YAP Pathway Suppression

Influenza infection suppressed the Hippo/YAP mechanosensing pathway in the lung (Figure 7E), with six genes downregulated including *Yap1* (−0.64), *Wwtr1*/TAZ (−0.42), *Lats1* (−0.31), *Lats2* (−0.84), *Amotl1* (−0.72), and *Amotl2* (−1.04). YAP1 and TAZ are transcriptional co-activators that sense mechanical stimuli including shear stress and matrix stiffness, driving tissue repair and alveolar epithelial regeneration after injury (92, 93). Their suppression removes the mechanosensing machinery required for alveolar repair, compounding the alveolar identity collapse and vascular repair failure described in section 3.4. Hippo/YAP suppression in the lung was confirmed by qRT-PCR, with *Yap1* showing significant downregulation consistent with the RNA-seq findings (Figure 7F).

In the aorta, seven Hippo pathway genes were similarly downregulated(94), including *Yap1* (−0.35), *Wwtr1*/TAZ (−0.22), *Lats2* (−0.40), *Amotl1* (−0.28), *Amotl2* (−0.56), *Ajuba* (−0.35), and *Ankrd1* (−1.93) (Figure 7G). YAP1 suppression in vascular smooth muscle cells has been directly linked to phenotypic switching toward a synthetic, pro-inflammatory state (93, 95), providing a mechanistic connection between Hippo pathway loss and the SMC dedifferentiation described in section 3.12. These data suggest that influenza infection suppresses the Hippo/YAP mechanosensing pathway in both tissues simultaneously, abolishing the alveolar repair response in the lung while promoting SMC dedifferentiation in the aorta, connecting a shared transcriptional vulnerability to two distinct tissue-specific mechanisms of structural failure. Hippo/YAP suppression in the aorta was confirmed by qRT-PCR, with *Yap1* showing significant downregulation consistent with the RNA-seq findings (Figure 7H).

### 3.16 Bone Marrow-Derived Macrophage Validation of In Vivo Transcriptional Programs

Macrophages are central mediators of both influenza-driven pulmonary injury and atherosclerotic plaque destabilization, making them a critical cellular link between the pulmonary and vascular transcriptional responses identified in this study. To validate the macrophage contribution to the in vivo transcriptional response, BMDMs were generated from *Apoe*−/− mice and treated with influenza A/PR/8/34 at MOI 1 or PBS for 24 hours. Genes representing seven distinct macrophage inflammatory mechanisms identified in the in vivo transcriptomes were selected for validation: innate viral sensing (*Zbp1*), inflammasome activation (*Pycard*), DAMP amplification (*S100a8*, *Trem1*, *Fpr2*), M1 polarization (*Nos2*), prostaglandin-mediated inflammatory resolution (*Hpgd*), and TGF-β quiescence signaling (*Tgfb1*) (71, 96). Influenza infection induced significant upregulation of *Zbp1*, *Nos2*, *Pycard*, *S100a8*, *Trem1*, and *Fpr2*, and significant downregulation of *Hpgd* and *Tgfb1* (Figure 8A–8H), confirming that all seven mechanisms are recapitulated in isolated macrophages independent of the tissue microenvironment.

**Figure 8.**
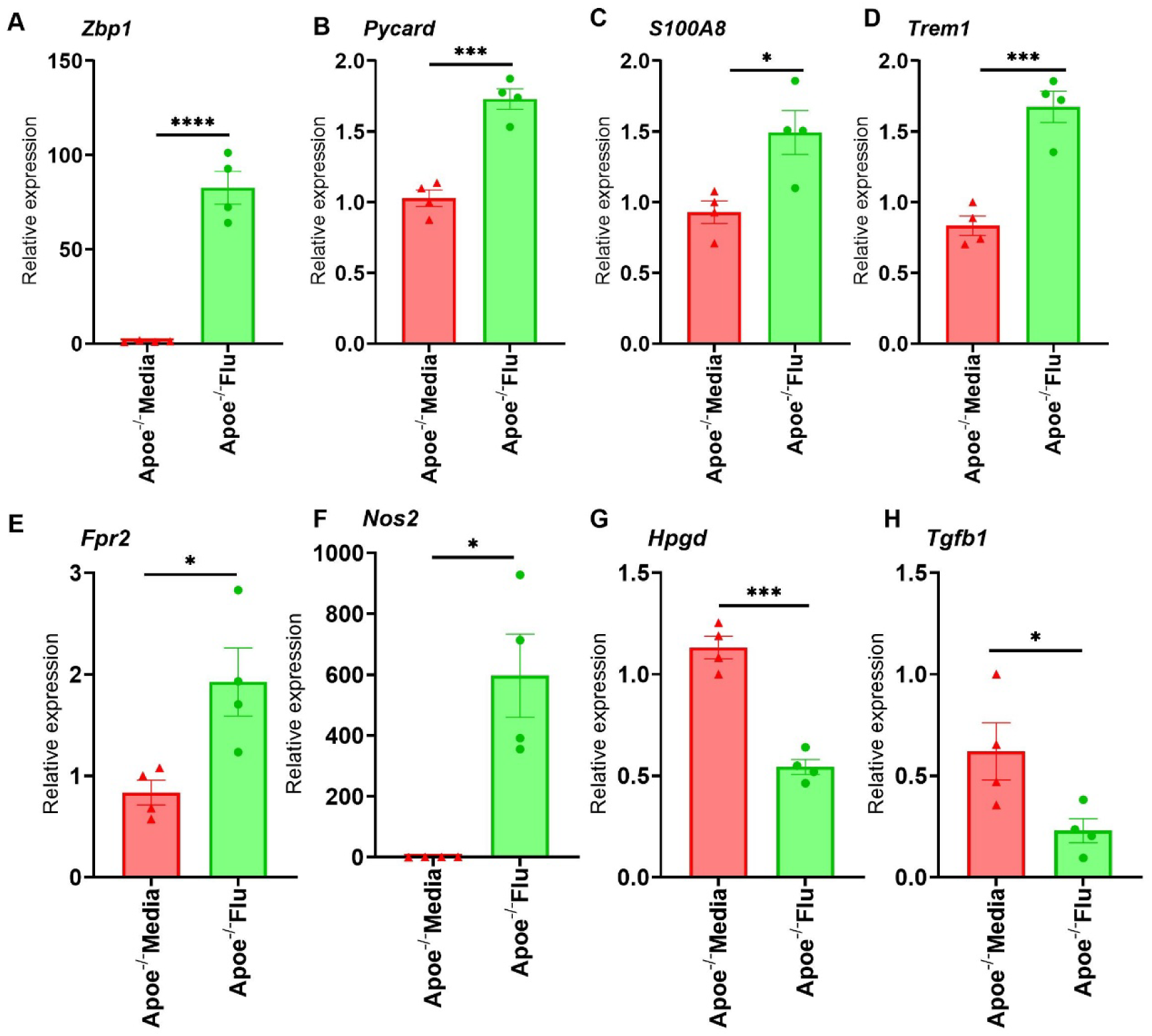
***Apoe*−/− macrophage-driven inflammatory response to influenza infection.** Bone marrow-derived macrophages from *Apoe*−/− mice were treated with PBS or influenza A/PR/8/34 at MOI 1 for 24 hours in DMEM. Relative expression of *Zbp1* (A), *Pycard* (B), *S100a8* (C), *Trem1* (D), *Fpr2* (E), *Nos2* (F), *Hpgd* (G), and *Tgfb1* (H) measured by qRT-PCR (n=4 per group). Data are presented as mean ± SEM. *p < 0.05 by unpaired two-tailed t-test. BMDM, bone marrow-derived macrophage; MOI, multiplicity of infection.

*Zbp1* upregulation (Figure 7A) suggests that Z-RNA sensing and downstream necroptotic and inflammasome signaling are activated in macrophages by influenza infection, consistent with its strong induction in the lung transcriptome (+2.02) (15). *Pycard* induction (Figure 7B) suggests macrophage inflammasome priming, consistent with the NLRP3 activation predicted by *Scd1* suppression in the aortic transcriptome (97). The induction of *S100a8* (Figure 7C), *Trem1* (Figure 7D), and *Fpr2* (Figure 7E) directly mirrors their upregulation in the aortic transcriptome (*S100a8* +1.63, *Trem1* +1.97, *Fpr2* +1.94), confirming that DAMP sensing and myeloid activation at the aortic wall have a macrophage-driven component independent of the tissue environment (98). *Nos2* induction (Figure 7F) indicates M1 macrophage polarization, consistent with the pro-inflammatory activation state identified in both lung and aortic transcriptomes (99). Downregulation of *Hpgd* (Figure 7G) suggests macrophage-driven failure of prostaglandin catabolism, creating a state of non-resolving inflammation through PGE2 accumulation that sustains macrophage activation independent of systemic cytokine signals (100). Downregulation of *Tgfb1* (Figure 7H) indicates that influenza-infected macrophages lose their TGF-β quiescence signal in a cell-intrinsic manner, consistent with *Tgfb2* and *Tgfb3* suppression observed at the aortic wall and the reduced anti-inflammatory macrophage populations documented in this model (26).

Together these findings demonstrate that influenza infection programs *Apoe*−/− macrophages into a sustained pro-inflammatory state through seven converging mechanisms, providing a cellular basis for the macrophage-driven components of the in vivo transcriptional response and supporting a role for macrophage programming in the persistence of cardiovascular risk during influenza recovery.

## 4. Discussion

This study provides the first genome-wide transcriptomic characterization of influenza-exacerbated atherosclerosis at the recovery phase, identifying 6,828 differentially expressed genes across the lung and aorta of HFD-fed *Apoe*−/− mice at day 15 post-infection. Three converging mechanisms were identified by which pulmonary influenza infection drives atherosclerotic plaque destabilization: a pulmonary inflammatory axis delivering activated monocytes, T cells, and acute phase proteins to the aortic wall, a metabolic axis in which convergent bioenergetic failure and NLRP3 inflammasome priming drive necrotic core expansion, and a structural axis in which SMC dedifferentiation and Hippo/YAP mechanosensing suppression drive fibrous cap thinning (6–8). Together these mechanisms provide a molecular basis for the epidemiological observation that cardiovascular mortality risk remains elevated for weeks after resolution of acute influenza symptoms.

The pulmonary transcriptional response reveals a self-amplifying inflammatory program extending well beyond classical antiviral immunity. The simultaneous induction of all four CCR2 ligands (*Ccl2*, *Ccl7*, *Ccl8*, *Ccl12*) represents the complete Ly6C-high monocyte mobilization program, generating the circulating monocyte pool that directly seeds aortic plaque macrophages (31, 32). The concurrent Th1, Th17, and exhaustion T-cell programs reflect the dual demand of viral clearance and tissue damage response at day 15, consistent with the CD4^+^RORγt^+^ and CD8^+^ T-cell expansions documented by flow cytometry in this model (14). The strong activation of the complement system, anchored by *C1qa*, *C1qb*, *C1qc*, and *C3ar1* alongside *Cfh*, alongside simultaneous efferocytosis failure through *Cd36*, *Tyro3*, and *Gas6* loss, establishes conditions for sustained pulmonary inflammation (101). The breadth of metabolic suppression, encompassing OXPHOS, glycolysis, cholesterol biosynthesis, and the transsulfuration pathway, extends beyond previously reported influenza-driven metabolic changes and provides a transcriptional explanation for why inflammatory resolution fails despite reduced viral burden. *Cbs* and *Cth* suppression, causing homocysteine accumulation and glutathione depletion, represent previously unrecognized metabolic vulnerabilities connecting pulmonary influenza infection to endothelial dysfunction and oxidative plaque injury (62).

The aortic transcriptome reveals a vascular response that directly recapitulates the cellular events of plaque destabilization. The simultaneous upregulation of eight MMPs in the lung, with *Mmp12* showing the strongest macrophage-specific induction, provides the enzymatic basis for fibrous cap degradation when these activated macrophages and their secreted products reach the aortic wall (102, 103). At the aortic wall, *Trem1* and *Fpr1/Fpr2* upregulation amplifies macrophage DAMP sensing in response to mitochondrial damage products released from necrotic plaque cells, while the class-switched humoral response confirmed by *Ighg1* (+7.13) indicates antigen-experienced B cells or plasma cells at the aortic wall, providing complement activation and Fc receptor-mediated macrophage stimulation (104, 105). Cytotoxic T-cell engagement confirmed by *Gzmb* (+2.85), combined with *Ccr7* downregulation trapping effector T cells within the aortic tissue and *Ccl17* loss removing the Treg recruitment signal, creates an unregulated inflammatory microenvironment that mirrors the immune composition of vulnerable human coronary plaques (106). The metabolic axis operating in the lung is mirrored in the aorta through convergent bioenergetic failure involving *Hk2*, *Aldoa*, and *Acaca*, with *Scd1* suppression in both compartments priming the NLRP3 inflammasome and amplifying the inflammatory program across tissues (96, 107). SMC dedifferentiation through *Myocd* loss, ferroptosis susceptibility through *Gpx4* suppression, and NLRP3 priming through *Scd1* suppression converge on fibrous cap thinning and necrotic core expansion, the two structural changes most predictive of plaque rupture (78). The stronger induction of *Apol11b* in the aorta than the lung indicates local IFN amplification beyond systemic spillover, consistent with direct endothelial cell infection demonstrated in this model (14) and identifies the aortic wall as an active rather than passive participant in the antiviral response. The acute phase response is particularly striking in its magnitude. *Saa3* was among the most strongly induced genes in the entire dataset (+6.36 log₂FC), expected to fundamentally remodel circulating HDL toward a pro-inflammatory, cholesterol-poor phenotype, promoting macrophage foam cell formation and endothelial adhesion molecule expression in the aortic wall (49, 103). Combined with *A2m* upregulation in both tissues, this sustained acute phase signal provides a prolonged atherogenic stimulus that outlasts the period of active viral replication and immune cell infiltration (108).

Three findings in this dataset are novel and warrant specific discussion. First, the ectopic upregulation of seven lung-identity genes in the aorta, precisely mirrored by their suppression in the lung, identifies a novel cross-organ communication mechanism. The simultaneous suppression of *Nkx2-1*, *S*ftpa1, *Sftpb*, *Sftpc*, *Scgb1a1*, *Scgb3a2*, and *Foxj1* in the lung alongside their induction in the aorta is consistent with lung-derived extracellular vesicle transfer or circulating transcripts delivering alveolar signals to the vascular wall during influenza-driven alveolar damage, warranting experimental investigation (109). Second, co-suppression of the molecular circadian clock and Hippo/YAP mechanosensing pathway across both tissues simultaneously identifies systemic transcriptional vulnerabilities connecting pulmonary infection to vascular structural failure through mechanisms independent of classical immune activation (90, 94, 95, 110). Circadian disruption through *Nr1d1* and *Arntl* loss removes the temporal restraint on NF-κB vascular inflammatory signaling, while YAP1 suppression in aortic SMCs drives phenotypic switching toward a synthetic state that thins the fibrous cap, providing a mechanistic link between influenza-driven mechanosensing loss and plaque rupture risk (91, 94). Third, influenza infection of *Apoe*−/− BMDMs induced a macrophage-driven inflammatory program encompassing innate viral sensing through *Zbp1*, inflammasome activation through *Pycard*, DAMP amplification through *S100a8*, *Trem1*, and *Fpr2*, M1 polarization through *Nos2*, suppression of inflammatory resolution through *Hpgd*, and loss of TGF-β quiescence signaling through *Tgfb1*(15, 104). The concordance between *S100a8*, *Trem1*, and *Fpr2* induction in BMDMs and their upregulation in the aortic transcriptome suggests that macrophage DAMP sensing at the aortic wall has a macrophage-driven component independent of the systemic inflammatory environment. Downregulation of *Tgfb1* in influenza-infected *Apoe*−/− macrophages indicates that macrophages lose their anti-inflammatory TGF-β quiescence signal in a cell-autonomous manner, consistent with *Tgfb2* and *Tgfb3* suppression at the aortic wall (111).These findings provide a cellular mechanism for the macrophage-driven components of both the lung and aortic transcriptional responses and identify macrophage programming as a contributor to the disproportionate cardiovascular risk of influenza in individuals with pre-existing atherosclerosis.

These findings have direct clinical implications specific to the three mechanisms identified. The pulmonary inflammatory axis, driven by persistent CCR2 monocyte mobilization and CXCR3 chemokine signaling, identifies a therapeutic window in which CCR2 or CXCR3 antagonism during the recovery phase could reduce monocyte delivery to the aortic wall and lower plaque inflammation (31, 32). The metabolic axis, defined by *Scd1* suppression and NLRP3 priming across both tissues, suggests that NLRP3 inflammasome inhibitors such as colchicine, already showing cardiovascular benefit in clinical trials, may be particularly effective in the post-influenza recovery period (112, 113). The structural axis, connecting *Myocd* loss, Hippo/YAP suppression, and macrophage-driven *Tgfb1* loss to fibrous cap thinning, raises the possibility that YAP pathway activators or TGF-β supplementation could preserve both SMC contractile identity and macrophage anti-inflammatory tone during influenza recovery (90, 95). Annual influenza vaccination, already associated with reduced cardiovascular events, likely attenuates these transcriptional programs upstream of all three mechanisms, and testing in vaccinated *Apoe*−/− mice would determine which axis is most effectively targeted by vaccination(114, 115).

### Limitations of Study

Several limitations should be noted. Bulk RNA-seq does not resolve cell-type-specific contributions and single-cell RNA-seq would be required to attribute pathway responses to specific cell populations. This analysis represents a single time point, and longitudinal sampling would define temporal dynamics from acute infection through recovery. While *Il17a/f* and *Rorc* were not detected in the bulk RNA-seq data, likely due to tissue dilution, upstream polarizing signals and downstream effectors provide transcriptional support for Th17 activity. Finally, delineating the relative contributions of lung-derived systemic signals and local plaque amplification would require adoptive transfer experiments or intravital imaging.

## 5. Conclusions

RNA sequencing of aortic and lung tissues from HFD-fed *Apoe*−/− mice reveals a lung-to-aorta inflammatory and metabolic axis driving atherosclerotic plaque destabilization through three converging mechanisms. In the lung, the IFN-γ-driven CXCR3 chemokine axis, comprehensive T-cell activation spanning Th1, Th17, and exhaustion programs, MMP-mediated ECM remodeling, alveolar identity collapse, and broad metabolic dysregulation collectively establish the lung as the transcriptional engine of cardiovascular risk amplification. In the aorta, convergent immune activation, bioenergetic failure, and SMC dedifferentiation simultaneously thin the fibrous cap and expand the necrotic core. The ectopic upregulation of lung-identity genes in the aorta, precisely mirrored by their suppression in the lung, identifies a novel cross-organ communication mechanism, and circadian clock disruption with Hippo/YAP pathway suppression shared across both tissues represent additional systemic vulnerabilities connecting pulmonary infection to vascular structural compromise. Influenza infection of *Apoe*−/− macrophages induced a macrophage-driven inflammatory program encompassing innate viral sensing, inflammasome activation, and DAMP amplification, identifying a cellular basis for the disproportionate cardiovascular risk of influenza in patients with pre-existing atherosclerosis. These transcriptional programs provide a mechanistic framework for evaluating anti-inflammatory and vaccination strategies targeting influenza-associated cardiovascular mortality.

## Author contributions

Conceptualization, R.G, L.D, and N.J.C; methodology, L.D, J.C.N, N.J.C, D.R and R.G; software, D.R and R.G; validation, L.D, J.C.N, N.J.C and R.G; formal analysis, R.G and D.R; investigation, L.D, J.C.N, N.J.C, D.R and R.G; resources, R.G and J.F.A; data curation, RG and D.R; writing – original draft, R.G; writing – review and editing, R.G and J.F.A; visualization, RG; supervision, RG; project administration, R.G; funding acquisition, R.G and J.F.A.

## Acknowledgements

The authors thank the Bioinformatics Core in the Department of Pediatrics, University of Pittsburgh for RNA-seq data processing and differential expression analysis. The authors acknowledge the use of Anthropic Claude (accessed July and August 2026) to polish the original draft of the manuscript. All AI-assisted text was reviewed and revised by the authors to ensure accuracy and clarity of meaning.

## Data availability statement

The RNA-seq data generated in this study will be deposited in the NCBI Gene Expression Omnibus (GEO) and will be publicly available as of the date of publication. The accession number will be provided upon acceptance. All original data reported in this paper will be shared by the lead contact upon request. This paper does not report original code. Any additional information required to reanalyze the data reported in this paper is available from the corresponding author upon request.

---

**Supplementary Table 1.**
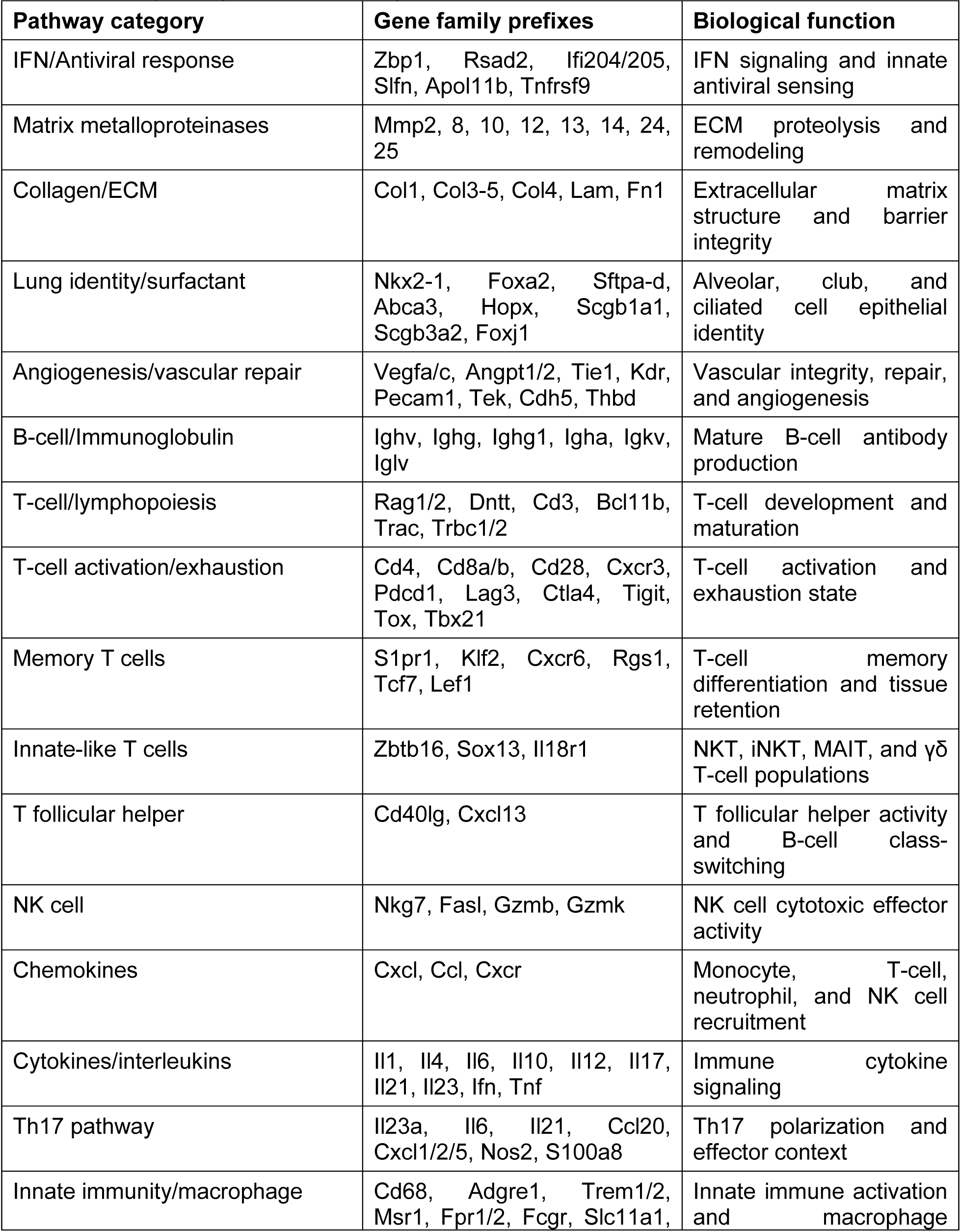
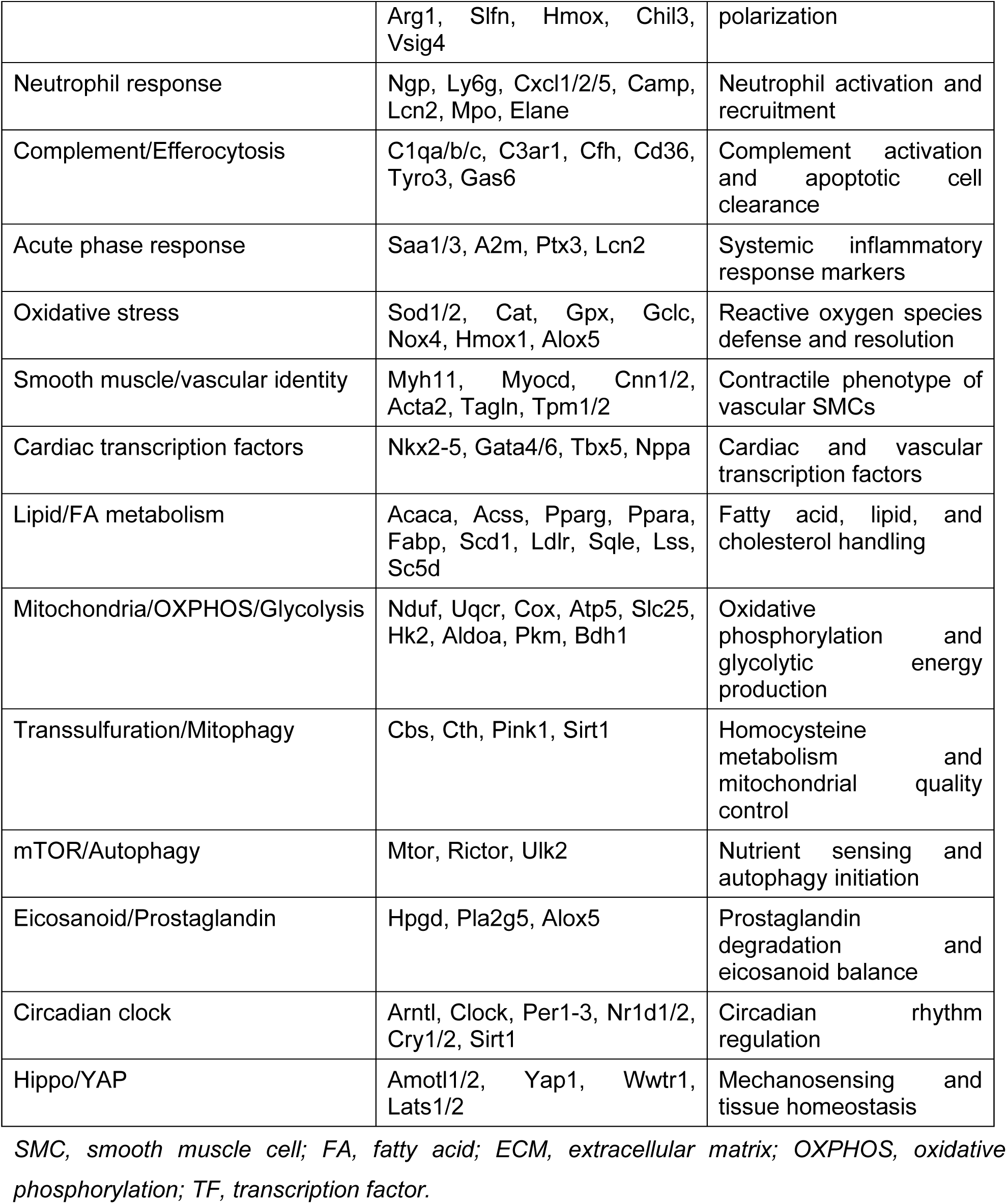
Gene pathway categories and biological functions used for heatmap classification.

**Supplementary Table 2.**
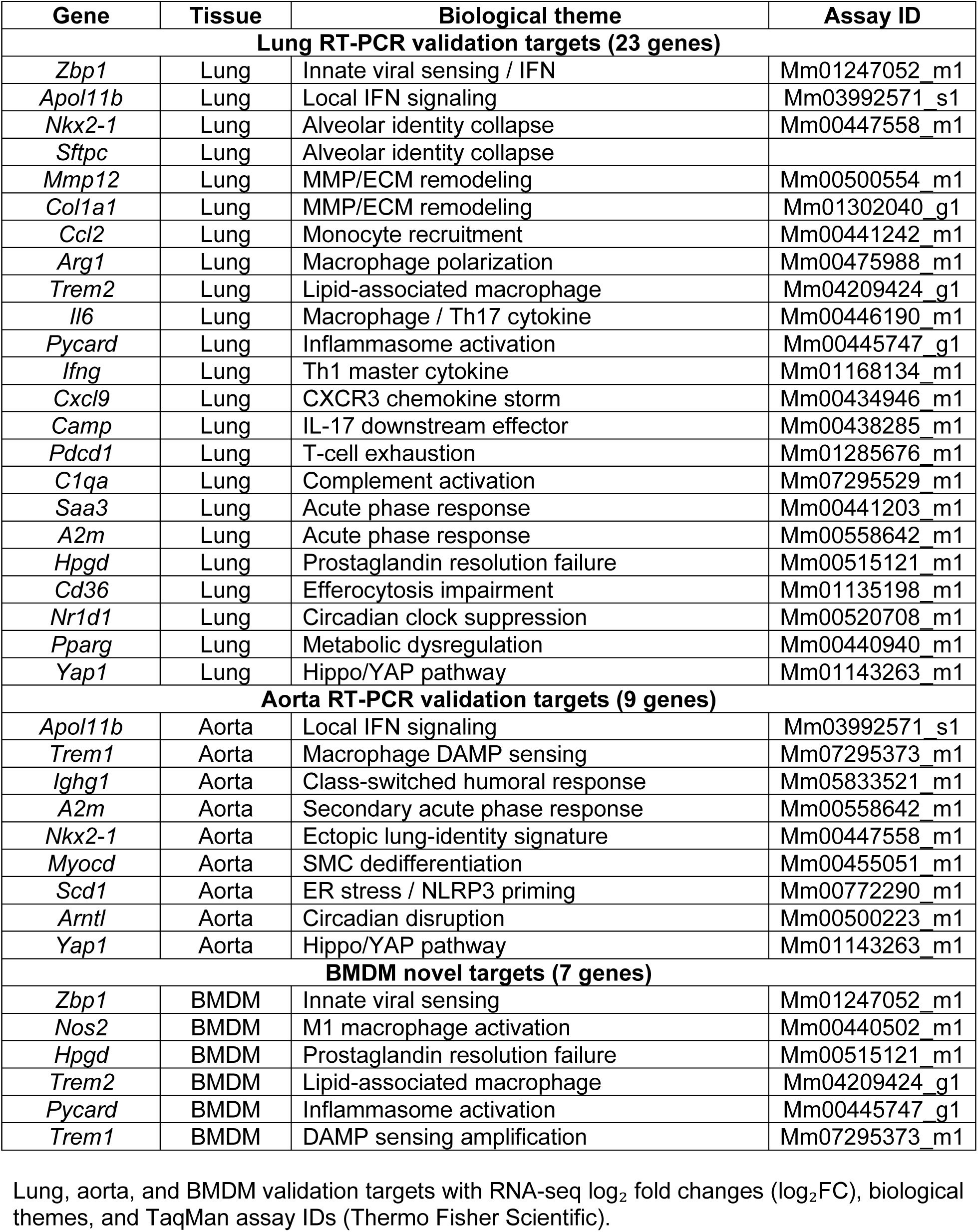
Gene expression validation targets for RNA-seq findings in lung, aorta, and BMDMs.

