## Supplemental File 1 for "Genome-Wide Transcriptomics Uncovers a Lung-to-Aorta Inflammatory Program Driving Influenza-Induced Atherosclerosis Exacerbation"

**Supplementary Table 1.**

**Gene pathway categories and biological functions used for heatmap classification**

| Pathway category | Gene family prefixes | Biological function |
|---|---|---|
| IFN/Antiviral response | Zbp1, Rsad2, Ifi204/205, Slfn, Apol11b, Tnfrsf9 | IFN signaling and innate antiviral sensing |
| Matrix metalloproteinases | Mmp2, 8, 10, 12, 13, 14, 24, 25 | ECM proteolysis and remodeling |
| Collagen/ECM | Col1, Col3-5, Col4, Lam, Fn1 | Extracellular matrix structure and barrier integrity |
| Lung identity/surfactant | Nkx2-1, Foxa2, Sftpa-d, Ager, Abca3, Hopx, Scgb1a1, Scgb3a2, Foxj1 | Alveolar, club, and ciliated cell epithelial identity |
| Angiogenesis/vascular repair | Vegfa/c, Angpt1/2, Tie1, Kdr, Pecam1, Tek, Cdh5, Thbd | Vascular integrity, repair, and angiogenesis |
| B-cell/Immunoglobulin | Ighv, Ighg, Ighg1, Igha, Igkv, Iglv | Mature B-cell antibody production |
| T-cell/lymphopoiesis | Rag1/2, Dntt, Cd3, Bcl11b, Trac, Trbc1/2 | T-cell development and maturation |
| T-cell activation/exhaustion | Cd4, Cd8a/b, Cd28, Cxcr3, Pdcd1, Lag3, Ctla4, Tigit, Tox, Tbx21, Icos | T-cell activation and exhaustion state |
| Memory T cells | S1pr1, Klf2, Cxcr6, Rgs1, Tcf7, Lef1 | T-cell memory differentiation and tissue retention |
| Innate-like T cells | Zbtb16, Sox13, Il18r1 | NKT, iNKT, MAIT, and γδ T-cell populations |
| T follicular helper | Icos, Cd40lg, Cxcl13 | T follicular helper activity and B-cell class-switching |
| NK cell | Nkg7, Fasl, Gzmb, Gzmk | NK cell cytotoxic effector activity |
| Chemokines | Cxcl, Ccl, Cxcr | Monocyte, T-cell, neutrophil, and NK cell recruitment |
| Cytokines/interleukins | Il1, Il4, Il6, Il10, Il12, Il17, Il21, Il23, Il27ra, Il17rd, Ifn, Tnf | Immune cytokine signaling |
| Th17 pathway | Il23a, Il6, Il21, Ccl20, Cxcl1/2/5, Nos2, S100a8, Il27ra, Traf6 | Th17 polarization and effector context |

| | | |
|---|---|---|
| Innate immunity/macrophage | Cd68, Adgre1, Trem1/2, Msr1, Fpr1/2, Fcgr, Slc11a1, Arg1, Slfn, Hmox, Chil3, Vsig4 | Innate immune activation and macrophage polarization |
| Neutrophil response | Ngp, Ly6g, Cxcl1/2/5, Camp, Lcn2, Mpo, Elane | Neutrophil activation and recruitment |
| Complement/Efferocytosis | C1qa/b/c, C3ar1, Cfh, Cd36, Tyro3, Gas6 | Complement activation and apoptotic cell clearance |
| Acute phase response | Saa1/3, A2m, Ptx3, Lcn2 | Systemic inflammatory response markers |
| Oxidative stress | Sod1/2, Cat, Gpx, Gclc, Nox4, Hmox1, Alox5 | Reactive oxygen species defense and resolution |
| Smooth muscle/vascular identity | Myh11, Myocd, Cnn1/2, Acta2, Tagln, Tpm1/2 | Contractile phenotype of vascular SMCs |
| Cardiac transcription factors | Nkx2-5, Gata4/6, Tbx5, Nppa | Cardiac and vascular transcription factors |
| Lipid/FA metabolism | Acaca, Acss, Pparg, Ppara, Fabp, Scd1, Ldlr, Sqle, Lss, Sc5d | Fatty acid, lipid, and cholesterol handling |
| Mitochondria/OXPHOS/Glycolysis | Nduf, Uqcr, Cox, Atp5, Slc25, Hk2, Aldoa, Pkm, Bdh1 | Oxidative phosphorylation and glycolytic energy production |
| Transsulfuration/Mitophagy | Cbs, Cth, Pink1, Sirt1 | Homocysteine metabolism and mitochondrial quality control |
| mTOR/Autophagy | Mtor, Rictor, Ulk2 | Nutrient sensing and autophagy initiation |
| Eicosanoid/Prostaglandin | Hpgd, Pla2g5, Alox5 | Prostaglandin degradation and eicosanoid balance |
| Circadian clock | Arntl, Clock, Per1-3, Nr1d1/2, Cry1/2, Sirt1 | Circadian rhythm regulation |
| Hippo/YAP | Amotl1/2, Yap1, Wwtr1, Lats1/2 | Mechanosensing and tissue homeostasis |

*SMC, smooth muscle cell; FA, fatty acid; ECM, extracellular matrix; OXPHOS, oxidative phosphorylation; TF, transcription factor.*

**Supplementary Table 2**

**Gene expression validation targets for RNA-seq findings in lung, aorta, and BMDMs**

| Gene | Tissue | Biological theme | Assay ID |
|---|---|---|---|
| **Lung RT-PCR validation targets (23 genes)** | | | |
| *Zbp1* | Lung | Innate viral sensing / IFN | Mm01247052_m1 |
| *Apol11b* | Lung | Local IFN signaling | Mm03992571_s1 |
| *Nkx2-1* | Lung | Alveolar identity collapse | Mm00447558_m1 |
| *Sftpc* | Lung | Alveolar identity collapse | |
| *Mmp12* | Lung | MMP/ECM remodeling | Mm00500554_m1 |
| *Col1a1* | Lung | MMP/ECM remodeling | Mm01302040_g1 |
| *Ccl2* | Lung | Monocyte recruitment | Mm00441242_m1 |
| *Arg1* | Lung | Macrophage polarization | Mm00475988_m1 |
| *Trem2* | Lung | Lipid-associated macrophage | Mm04209424_g1 |
| *Il6* | Lung | Macrophage / Th17 cytokine | Mm00446190_m1 |
| *Pycard* | Lung | Inflammasome activation | Mm00445747_g1 |
| *Ifng* | Lung | Th1 master cytokine | Mm01168134_m1 |
| *Cxcl9* | Lung | CXCR3 chemokine storm | Mm00434946_m1 |
| *Camp* | Lung | IL-17 downstream effector | Mm00438285_m1 |
| *Pdcd1* | Lung | T-cell exhaustion | Mm01285676_m1 |
| *C1qa* | Lung | Complement activation | Mm07295529_m1 |
| *Saa3* | Lung | Acute phase response | Mm00441203_m1 |
| *A2m* | Lung | Acute phase response | Mm00558642_m1 |
| *Hpgd* | Lung | Prostaglandin resolution failure | Mm00515121_m1 |
| *Cd36* | Lung | Efferocytosis impairment | Mm01135198_m1 |
| *Nr1d1* | Lung | Circadian clock suppression | Mm00520708_m1 |
| *Pparg* | Lung | Metabolic dysregulation | Mm00440940_m1 |
| *Yap1* | Lung | Hippo/YAP pathway | Mm01143263_m1 |
| **Aorta RT-PCR validation targets (9 genes)** | | | |
| *Apol11b* | Aorta | Local IFN signaling | Mm03992571_s1 |
| *Trem1* | Aorta | Macrophage DAMP sensing | Mm07295373_m1 |
| *Ighg1* | Aorta | Class-switched humoral response | Mm05833521_m1 |
| *A2m* | Aorta | Secondary acute phase response | Mm00558642_m1 |
| *Nkx2-1* | Aorta | Ectopic lung-identity signature | Mm00447558_m1 |
| *Myocd* | Aorta | SMC dedifferentiation | Mm00455051_m1 |
| *Scd1* | Aorta | ER stress / NLRP3 priming | Mm00772290_m1 |
| *Arntl* | Aorta | Circadian disruption | Mm00500223_m1 |
| *Yap1* | Aorta | Hippo/YAP pathway | Mm01143263_m1 |
| **BMDM novel targets (7 genes)** | | | |
| *Zbp1* | BMDM | Innate viral sensing | Mm01247052_m1 |
| *Nos2* | BMDM | M1 macrophage activation | Mm00440502_m1 |
| *Hpgd* | BMDM | Prostaglandin resolution failure | Mm00515121_m1 |
| *Trem2* | BMDM | Lipid-associated macrophage | Mm04209424_g1 |
| *Pycard* | BMDM | Inflammasome activation | Mm00445747_g1 |
| *Trem1* | BMDM | DAMP sensing amplification | Mm07295373_m1 |

Lung, aorta, and BMDM validation targets with RNA-seq $\log_2$ fold changes ($\log_2$FC), biological themes, and TaqMan assay IDs (Thermo Fisher Scientific).
